# Flavonoid metabolism and its different physiological roles in plant cold acclimation

**DOI:** 10.1101/2024.06.13.598861

**Authors:** Anastasia Kitashova, Martin Lehmann, Serena Schwenkert, Dario Leister, Thomas Nägele

## Abstract

Flavonoids represent a diverse group of plant secondary metabolites which are also discussed in context of dietary health and inflammatory response. Numerous studies have revealed that flavonoids play a central role in plant acclimation to abiotic factors like low temperature or high light, but their structural and functional diversity frequently prevents a detailed mechanistic understanding. Further complexity in analysing flavonoid metabolism arises from the different subcellular compartments which are involved in biosynthesis and storage. In the present study, non-aqueous fractionation of *Arabidopsis* leaf tissue was combined with metabolomics and proteomics analysis to reveal subcellular regulation during cold acclimation in two flavonoid-deficient mutants. During the first three days of a two-week cold acclimation period, flavonoid deficiency was observed to affect pyruvate, citrate and glutamate metabolism which indicated a role in stabilising C/N metabolism and photosynthesis. Also, tetrahydrofolate metabolism was found to be affected which had significant effects on regulation of the proteome of the photorespiratory pathway. In the late stage of cold acclimation, flavonoid deficiency was found to affect protein stability, folding and proteasomal degradation which resulted in a significant decrease of total protein amounts in both mutants. In summary, these findings suggest that flavonoid metabolism plays different roles in early and late stages of plant cold acclimation and significantly contributes to establishing a new protein homeostasis in a changing environment.

## Introduction

Plant metabolism comprises numerous metabolic compounds, belonging to diverse chemical substance classes, which are synthesized, degraded and interconverted enzymatically within comprehensive and highly interconnected metabolic networks. Central plant primary metabolites are carbohydrates which are direct products of photosynthetic CO_2_ fixation and, thus, represent substrates for all further anabolic metabolic pathways. Metabolism of carbohydrates is tightly linked to carboxylic and amino acids, which, all together, represent the building blocks for protein, lipid and nucleotide biosynthesis. Further, they also represent substrates for biosynthesis of diverse secondary compounds, e.g., alkaloids, glucosinolates or flavonoids, which have been shown to protect plants against environmental stressors, biotic and abiotic ones, by their specialised functions (Erb and Kliebenstein 2020).

In many plant species, exposure to low but non-freezing temperatures induces a process termed cold acclimation which increases their freezing tolerance. This process has been studied for decades and has revealed numerous molecular and physiological responses which are involved in its regulation (Steponkus 1984; Ristic and Ashworth 1993; Gilmour et al. 1998; Kaplan et al. 2007; Garcia-Molina et al. 2020). Adjustment of photosynthesis and carbohydrate metabolism is a central cold acclimation response to prevent or mitigate photoinhibition and a disbalance between primary and secondary photosynthetic reactions (Stitt and Hurry 2002). Sucrose metabolism has been found to play a crucial role in photosynthetic acclimation (Strand et al. 2003). It has been discussed that that adjustment of sucrose phosphate synthase (SPS) activity to low temperature prevents a limitation of triose phosphate/phosphate translocation between cytosol and chloroplasts which would result in a limitation of photosynthetic ATP biosynthesis (Nägele et al. 2012). Recently, we found evidence for a role of SPS activity in regulation of carbon fluxes between carbohydrate and flavonoid metabolism during cold acclimation (Kitashova et al. 2023). While metabolism of flavonoids is well known to be involved in many plant-environment interactions (Winkel-Shirley 2002), their physiological role in cold acclimation still remains elusive. Flavonoids are phenylpropanoids which are synthesised from phenylalanine and tyrosine through the shikimic acid pathway. For *Brassica napus*, phenylpropanoid deficiency has been found to result in lowered freezing tolerance and in a decreased cold acclimation capacity of photosynthesis (Solecka and Kacperska 2003). Also, for *Arabidopsis*, a significant impact of flavonoids and their metabolism on cold acclimation capacity and freezing tolerance has been described (Schulz et al. 2016, 2015). Under high light, recent findings suggested flavonoid metabolism to be tightly interconnected with chloroplast carbohydrate metabolism and the SnRK1-related signalling network (Zirngibl et al. 2023). Other studies have provided further evidence for diverse functional roles of flavonoids, comprising, e.g., DNA protection, protection against UV radiation or serving as signalling compounds to affect gene expression (Sarma and Sharma 1999; Naoumkina and Dixon 2008; Nakabayashi et al. 2014). These findings, together with observations made in many other studies (for an overview, see e.g., (Tohge et al. 2018)), provide strong evidence for a central role of flavonoids in plant stress response and cold acclimation.

Flavonoid accumulation is typically observed in the vacuole (for an overview, see e.g. (Pucker and Selmar 2022)). Also, flavonoids have been reported to be transported to other compartments, e.g., the nucleus and chloroplasts (Peer et al. 2001; Agati et al. 2007). Together with the structural diversity of flavonoids, also the diversity of subcellular localization challenges the analysis of their plant biochemical and physiological functions.

The core pathway of flavonoids biosynthesis is located in the cytosol, and comprises activities of chalcone synthase (CHS), chalcone isomerase (CHI), flavanone 3-hydroxylase (F3H), dihydroflavonol 4-reductase (DFR) and anthocyanidin synthase (ANS). Substrate for flavonoid biosynthesis is phenylalanine which is biosynthesised in plastids from erythrose 4-phosphate and phosphoenolpyruvate (Rippert et al. 2009). Erythrose 4-phosphate is a product of the Calvin-Benson-Bassham Cycle (CBBC) and, thus, directly links the shikimate pathway and subsequent flavonoid biosynthesis to photosynthetic CO_2_ fixation and carbohydrate biosynthesis. Under changing environmental conditions, e.g., low temperature, photosynthetic acclimation prevents a disbalance of electron transport, ATP-biosynthesis and enzymatic CO_2_ fixation. Photosynthetic acclimation is a comprehensive process which comprises and affects regulation of redox and ion homeostasis of the chloroplast, protein amounts of photosystems, metabolism and multicompartmental signalling networks (Gjindali and Johnson 2023). Under low temperature, enzymes of the CBBC have been found to be upregulated to compensate for the decrease of activity due to thermodynamic constraints (Strand et al. 1999; Stitt and Hurry 2002). Together with regulation of the central carbohydrate metabolism, e.g., the sucrose biosynthesis pathway, these are immediate cold-induced processes which can typically be observed during the initial hours and days of a cold acclimation period (Savitch et al. 2001; Nägele et al. 2011). Accumulation of flavonoids and other specialized metabolites typically follows these initial acclimation reactions of primary metabolism, resulting in peak values after several days and up to weeks of cold exposure (Doerfler et al. 2013; Kitashova et al. 2023).

The balance between energy absorbed through photochemistry, photosynthetic electron transport and interconverting metabolic processes, also termed as photostasis, plays a critical role for plant-environment interactions in general (Huner et al. 2003). Here, regeneration of NADP^+^ as electron acceptor and ADP as substrate for the ATP synthase reaction are essential to prevent photoinhibition and generation of reactive oxygen species. The multicompartmental pathway of photorespiration has been discussed to play a central role of balancing ATP and NADPH production and usage (for an overview, see e.g. (Timm and Hagemann 2020)). Further, photorespiration affects many other pathways due to its role in C1 metabolism, S-metabolism or C/N balance.

To better understand the role of flavonoids during cold acclimation in a pathway-specific context, the present study analysed metabolism in flavonoid deficient mutants on a compartmental level. Subcellular metabolomics and proteomics data suggest a tight linkage of photorespiration and flavonoid metabolism through tetrahydrofolate-related C1 metabolism. The data further suggest a central role of this metabolic network for amino acid metabolism and stabilisation of the cellular protein homeostasis after cold exposure.

## Materials and Methods

### Plant material and growth conditions

Plants of *Arabidopsis thaliana* accession Col-0 and homozygous T-DNA insertion lines *chs* (chalcone synthase, line SALK_020583C, locus AT5G13930), and *f3h* (flavanone 3-hydroxylase, line SALK_113904C, locus AT3G51240) were grown as described before (Kitashova et al. 2023). In summary, plants were grown on a 1:1 mixture of GS90 soil and vermiculite in a climate chamber under controlled short-day conditions (8 h/16 h light/dark; 100 μmol m^−2^ s^−1^; 22°C/16°C; 60% relative humidity). After 4 weeks, plants were shifted to a growth cabinet and grown further under long day conditions (16 h/8 h light/dark; 100 μmol m^−2^ s^−1^; 22°C/16°C; 60% relative humidity). After 2 weeks, plants were either (i) harvested at midday, that is, 8 h of light (0 days of cold acclimation) or (ii) transferred to a cold room for cold acclimation (16 h/8 h light/ dark; 90–100 μmol m^−2^ s^−1^; 4°C/4°C). Cold exposed plants were harvested after 3 or 14 days of acclimation at midday, that is, after 8 h of light. Each sample consisted of nine leaf rosettes which were immediately frozen in liquid nitrogen, ground to a fine powder and lyophilised.

### Non-aqueous subcellular fractionation

Non-aqueous subcellular fractionation (NAF) was performed according to (Fürtauer et al. 2016) with slight modifications. In brief, approximately 10 mg of lyophilised plant material was homogenised in tetrachloroethylene (ρ = 1.60 g*cm^-3^) with an ultrasonic homogenizer (Hielscher Ultrasonics UP200St, www.hielscher.com). After centrifugation with 20.000 g, the supernatant was transferred in another tube, and its density was reduced with heptane (ρ = 0.68 g*cm^-3^). The pellet was then resuspended in tetrachloroethylene. Sonication of the supernatant with newly adjusted density, centrifugation, separation of pellet and density adjustment of the supernatant were repeated several times until a gradient of 5 to 7 densities was obtained. Each of the resulting resuspended pellets was split in 2 sub-fractions and dried in the desiccator. The resulting sub-gradients were used for (i) marker enzyme quantification; and (ii) primary metabolite quantification with GC-MS/MS. Activities of marker enzymes, i.e. plastidial pyrophosphatase, cytosolic uridine 5′diphosphoglucose pyrophosphorylase and vacuolar acidic phosphatase, were determined photometrically following (Fürtauer et al. 2016). The correlation between marker enzymes and the metabolite data was done according to the (Fürtauer et al. 2016).

### Extraction of lipids and primary metabolites for mass spectrometry

Lipids and non-polar metabolites were extracted following the method described by (Hummel et al. 2011). In summary, approximately 5 mg of freeze-dried leaf material or NAF fractionation pellets were subjected to extraction using 1 ml of a pre-cooled (-20°C) mixture of methanol and methyl tert-butyl ether (MTBE) in a ratio of 3:1. This extraction solution included 10 µl corticosterone, 5 µl chloramphenicol, 1.25 µl ampicillin, 2.5 µl sorbitol-^13^C, and 5 µl ribitol, each at a concentration of 0.2 mg ml^−1^, serving as internal standards. The mixture was vigorously mixed until the tissue was fully suspended and then shaken for 10 min at 4°C, followed by 10 min of sonication on ice. Phase separation was initiated by adding 500 µl of a water:methanol (3:1) solution, followed by vigorous mixing and centrifugation at maximum speed using a Centrifuge 5417R (Eppendorf, www.eppendorf.com). For lipid analysis, 600-700 µl of the upper phase was collected, while 400-500 µl of the lower phase was divided into equal aliquots for GC-TOF-MS analysis (sample dependent). All samples were dried using a vacuum concentrator (Concentrator 5301; www.eppendorf.com) and subsequently stored at - 80°C until further analysis. To prevent oxidation, argon was added during storage.

### LC-MS: lipid analysis

For LC-MS analysis, the Dionex Ultimate 3000 UHPLC (Thermo Fisher Scientific, www.thermofisher.com) in combination with a timsTOF (Bruker, www.bruker.com) was used. The dry extract was resolved in acetonitrile:isopropanol (7:3) and injected on a C8 reversed phase column (Ultra C8 100×2.1 mm; Restek, www.restek.com) with 300 µl min^−1^ flow at 60°C. The solvents used are (A) water and (B) acetonitrile:isopropanol (7:3), both including 1% (v/v) ammonium acetate and 0.1% (v/v) acetic acid. The 26 min gradient started at 55% B, followed by a ramp to 99% B within 15 min. After a 5 min washing step at 99% B, the gradient was returned to 55% B and kept constant for 5 min equilibration.

MS detection was performed using an electrospray ionization (ESI) source, operating in positive mode. Nitrogen served as the dry gas, at 8 L min^−1^, 8 bar, and 200°C. The timsTOF mass spectra were recorded in MS and MSMS mode from 50 m/z to 1300 m/z with 40.000 resolution, 1 Hz scan speed, and 0.3 ppm mass accuracy. Compounds were annotated in a targeted approach using the specific mass (m/z) at retention time and the isotopic pattern. All data were acquired by Compass HyStar 4.1 and otofControl 6.2. The evaluation was performed by DataAnalysis 5.1, and MetaboScape 2021. All software tools were provided by Bruker. Raw values are normalized by internal standard and gram dry weight.

### GC-MS: primary metabolite analysis

For the derivatization process, the pellet was suspended in 10 μl of methoxyamine hydrochloride solution (20 mg ml^-1^ in pyridine) and incubated for 90 min at 40°C. Following this, 20 µl of BSTFA (N,O-Bis[trimethylsilyl]trifluoroacetamide) containing 2.5 μl of a retention time standard mixture comprising linear alkanes (n-decane, n-dodecane, n-pentadecane, n-nonadecane, n-docosane, n-octacosane, n-dotriacontane) was added, and the mixture was further incubated at 40°C for 45 min. Subsequently, one and two μl of each sample was injected into a GC–TOF–MS system (Pegasus HT, Leco, www.leco.com) utilizing splitless injection as well as split ratios of 10, 30, and 50 (sample dependent). Sample injection and processing were automated using an autosampler system (Combi PAL, CTC Analytics AG, www.ctc.ch). Helium was employed as the carrier gas at a constant flow rate of 1 ml min^-1^. Gas chromatographic separation was conducted on an Agilent GC (7890A, Agilent, www.agilent.com) equipped with a 30 m VF-5ms column coupled with a 10 m EZ-Guard column. The temperature of the split/splitless injector, transfer line, and ion source was maintained at 250°C. The initial oven temperature was set at 70°C and ramped up to 350 °C at a rate of 9°C min^-1^. Metabolites were fractionated and ionized by a 70 eV ion pulse. Mass spectra were acquired at a rate of 20 scans s^-1^ within an m/z range of 50–600. Chromatograms and mass spectra were analysed using ChromaTOF 4.72 and TagFinder 4.1 software (Luedemann et al. 2008).

### Protein extraction and analysis by LC-MS/MS

Protein extraction and preparation for the LC-MS/MS quantification was done as previously described with minor adjustments (Marino et al. 2019). In summary, proteins were extracted using 6 M guanidine-chlorine in 10 mM HEPES pH 7.8, supplemented with 1 tablet of protease inhibitor cocktail per 10 ml buffer (cOmplete™ Proteasehemmer-Cocktail, ©Roche) with an ultrasonic homogenizer (Hielscher Ultrasonics UP200St, www.hielscher.com). Then, the proteins were precipitated with methanol/chloroform in water, washed with methanol, and the supernatant was discarded, leaving protein-containing pellets. Dried pellets were resuspended in 6 M Urea / 2 M Thiourea in 50 mM HEPES pH 7.8 buffer at 37°C. Total protein content was quantified using the Pierce™ 660nm Protein Assay Kit (©Thermo Scientific™). Following quantification, 80 µg of protein was aliquoted, reduced with 10 mM DTT, and alkylated with 50 mM iodoacetamide. The proteins were subsequently digested overnight with Trypsin at 37°C. Acidified with formic acid samples were purified using home-made C18 stage tips with 80% acetonitrile and 0.5% formic acid solution. Finally, 1 µg of purified protein extract was used for the LC-MS/MS analysis.

LC-MS/MS analysis was conducted as previously described with minor modifications, involving peptide separation over a 90 min linear gradient ranging from 5% to 80% (v/v) acetonitrile (ACN) (Espinoza-Corral et al. 2023). Raw data files were processed using MaxQuant software version 2.2.0.0 (Cox and Mann 2008), with peak lists searched against the *Arabidopsis* reference proteome from Uniprot (www.uniprot.org), employing default settings with ‘match-between-runs’ enabled. Protein quantification was achieved utilizing the label-free quantification algorithm (LFQ) (Cox et al. 2014). Subsequent analysis was conducted using Perseus version 2.0.9.0 (Tyanova et al. 2016). To refine the dataset, potential contaminants, proteins identified solely through site modification, and reverse hits were excluded. Only protein groups quantifiable by the LFQ algorithm in at least three out of four replicates under at least one condition were retained. LFQ intensities underwent log_2_ transformation, and missing values were imputed from a normal distribution within Perseus using standard settings.

### Statistics and data analysis

Dynamics of the subcellular metabolomes were analysed in MATLAB® (www.themathworks.com) using a method for time series analysis in context of biochemical network information (Nägele et al. 2016). In brief, sigmoidal Gompertz functions (Eq. 1) were fitted to experimental data (0➔3 days at 4°C, 3➔14 days at 4°C).

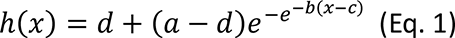

Gompertz functions describe growth rates with the slowest growth at the beginning of the period, i.e., reflecting the moment of cold exposure and thermodynamically affected enzymatic reaction rates. Derivatives of Gompertz functions represented metabolic functions, i.e., the summed rates of biosynthesis and degradation for each metabolite pool. Dynamics of metabolic functions were then compared to dynamics of substrate concentrations of each enzymatic reaction described by a metabolic network of subcellular carbohydrate metabolism, carboxylic acids and amino acids (Supplementary Figure 1). Dynamics were indicated by *ω*-functions (Eq. 2):

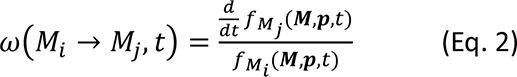

Here, *M_i_* denotes the substrate concentration of an enzymatic reaction and *M_j_* represents the product concentration. Metabolic functions of both substrate and product pools, *f*_*M*_*i*__ and *f*_*M*_*j*__, were derived from Gompertz functions as described before. To identify metabolic deregulation in both mutants, ratios of z-scaled omega values were built (Supplementary Figure 2, Supplementary Table 1).

Statistical analysis was performed with free software R version 4.2.2 (www.r-project.org) (R Core Team 2019) and RStudio version 2023.9.1.494 (www.rstudio.com). Figures were prepared using the RStudio or Python version 3.8.8 (www.python.org) and Jupyter Notebook version 6.3.0 (https://jupyter.org). The protein phylogeny analysis was performed using MUSCLE Multiple Sequence Alignment version 3.8, and pairwise sequence alignment was conducted using the EMBOSS Water tool (https://www.ebi.ac.uk). Pairwise protein structure alignment of proteins was performed using AlphaFold structures at https://www.rcsb.org. Ligand docking simulation was performed using the free software DockThor version 2.0 (https://dockthor.lncc.br/v2). The results from ligand docking were visualized using PyMOL version 2.5.7 (https://pymol.org). Evolutionary conservation profile of FOLD1 protein was performed using free software ConSurf (https://consurf.tau.ac.il). Differentially abundant proteins (DAP) were identified according to ANOVA significance test with Tukey HSD (Supplementary Table 2A). DAP GO term enrichment analysis (Biological Process) was performed using the “org.At.tair.db” package version 3.16.0 and “clusterProfiler” package version 4.6.2. GO terms were checked for redundancies with the most significant terms kept.

## Results

### Flavonoid deficiency affects regulation of citrate and glutamate metabolism during cold acclimation

Time series analysis of metabolomics data revealed that citrate and glutamate metabolism were consistently perturbed in both mutants, *chs* and *f3h*, during the full cold acclimation period of 14 days (Figure 1, Supplementary Table 1). In Col-0, cytosolic pyruvate, which is substrate for citrate biosynthesis via the pyruvate dehydrogenase reaction, was found to drop significantly during the first 3 days at 4°C and stabilised again until 14 days of cold acclimation. In both mutants, the cytosolic pyruvate amounts consistently decreased until 14 days at 4°C. In Col-0 and *chs*, citrate amounts decreased in all compartments until 3 days of cold before they consistently increased again. In *f3h*, cold-induced dynamics of citrate amounts were almost absent. Remarkably, both mutants showed higher variance of citrate amounts after 14 days at 4°C than Col-0. Furthermore, in *f3h*, a high variance of subcellular citrate amounts was observed across all time points of cold acclimation. Glutamate amounts showed an opposite trend in both mutants compared to Col-0. While cold acclimation resulted in increased glutamate amounts in Col-0 across all compartments, amounts decreased in both mutants until 14 days at 4°C, and the strongest decrease was observed in the plastidial fraction. Plastidial glutamine amounts in Col-0 decreased after 3 days of cold exposure which was followed by an increase after 14 days. In contrast, *chs* and *f3h* mutants demonstrated opposite trends: both mutants showed an increase in plastidial glutamine after 3 days, with no significant change in *chs* or with a decrease in *f3h* after 14 days at 4°C (Figure 1).

**Figure 1.**
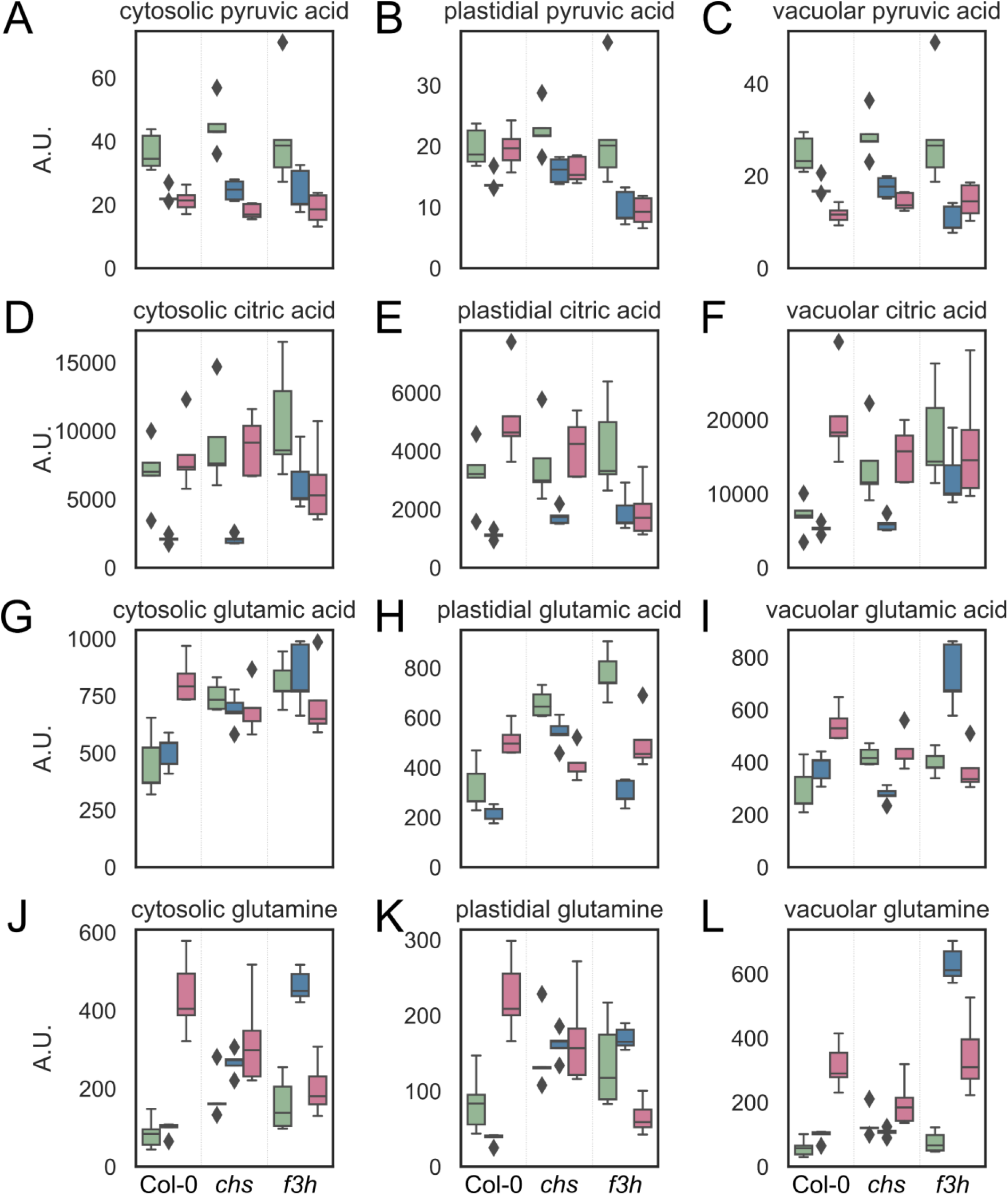
Cold-induced dynamics of the subcellular metabolome. Dynamics of (**A-C**) pyruvic acid, (**D-F**) citric acid, (**G-I**) glutamic acid and (**J-L**) glutamine in Col-0, *chs*, and *f3h* after 0, 3, and 14 days of cold acclimation. Green colour represents: 0 day; blue: 3 days; magenta: 14 days of cold acclimation; n=5. The list of significance levels (ANOVA with Tukey HSD) is provided in Supplementary Table 2B.

In addition to a perturbed pyruvate and citrate metabolism, also, amounts of pyruvate dehydrogenases (PDHs) and isocitrate dehydrogenases (ICDHs) were found to be affected in *chs* and *f3h* compared to Col-0 (Supplementary Figure 3). Remarkably, cytosolic ICDH (cICDH, At1G65930) significantly decreased in Col-0 until 3 days at 4°C and stabilised until 14 days. In both mutants, amounts of cICDH were significantly higher at 3 days and decreased to levels of Col-0 until 14 days. In contrast, mitochondrially located ICDHs showed similar dynamics across all genotypes. Interestingly, in the full data set, i.e., comprising all time points and genotypes, cICDH positively correlated with a plastidial pyruvate dehydrogenase (pPDH, AT1G01090) while it correlated negatively with a mitochondrial ICDH (AT2G17130) and a mitochondrial PDH (mPDH, AT1G59900; Figure 2).

**Figure 2.**
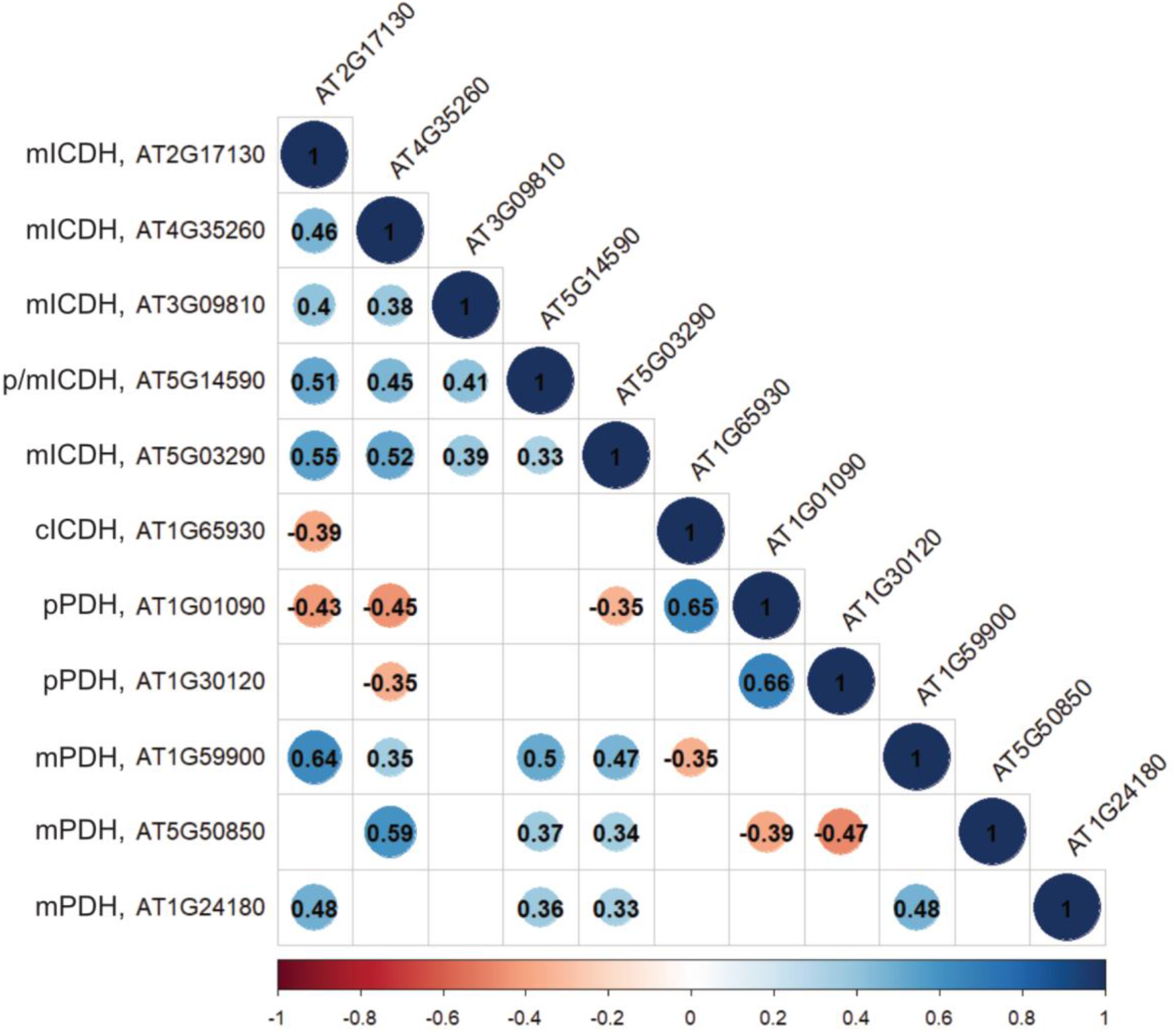
Pearson correlation of the subcellular proteome of pyruvate dehydrogenases and isocitrate dehydrogenases. Numbers show Pearson correlation coefficients; blank fields were not significant (p < 0.05). Blue: positive correlation; red: negative correlation. mICDH: mitochondrial isocitrate dehydrogenase; p/mICDH: plastidial/mitochondrial isocitrate dehydrogenase; cICDH: cytosolic isocitrate dehydrogenase; pPDH: plastidial pyruvate dehydrogenase; mPDH: mitochondrial pyruvate dehydrogenase.

### Flavonoid biosynthesis modulates photorespiration and amino acid metabolism dynamics through tetrahydrofolate-related C1 metabolism

Primary metabolites and lipids were quantified via chromatography coupled to mass spectrometry to reveal effects of flavonoid deficiencies on metabolic regulation during cold acclimation. Dimensionality reduction by principal component analysis (PCA) showed a general effect of cold acclimation on metabolomes and lipidomes (Figure 3), separating non-acclimated from cold-acclimated samples along PC1, while 3 days and 14 days of cold acclimation were separated along PC2. Loadings with highest contribution to separation along PC1 were carbohydrates (sucrose, maltose, glucose, fructose, raffinose), shikimic acid and glyceric acid. Along PC2, acetylserine, phosphoglyceric acid, malic acid, quinic acid, xylitol and galactonic acid contributed strongest to the separation (see Supplementary Table 3). Although the low temperature exposure had a dominant effect on sample separation (PC1), there was a subtle separation of Col-0 samples from metabolic mutants - which were either deficient in CHS (*chs*) or F3H (*f3h*) activity - after 3 days of cold. Thus, the absence of cytosolic flavonoid biosynthesis affected cold acclimation of primary metabolism and this effect was most pronounced during the early cold exposure period. An overview of growth phenotypes and PCA loadings, representing the contribution of individual metabolites and lipids to the observed separation, along with the components, is provided in the supplements (Supplementary Figure 4; Supplementary Table 3).

**Figure 3.**
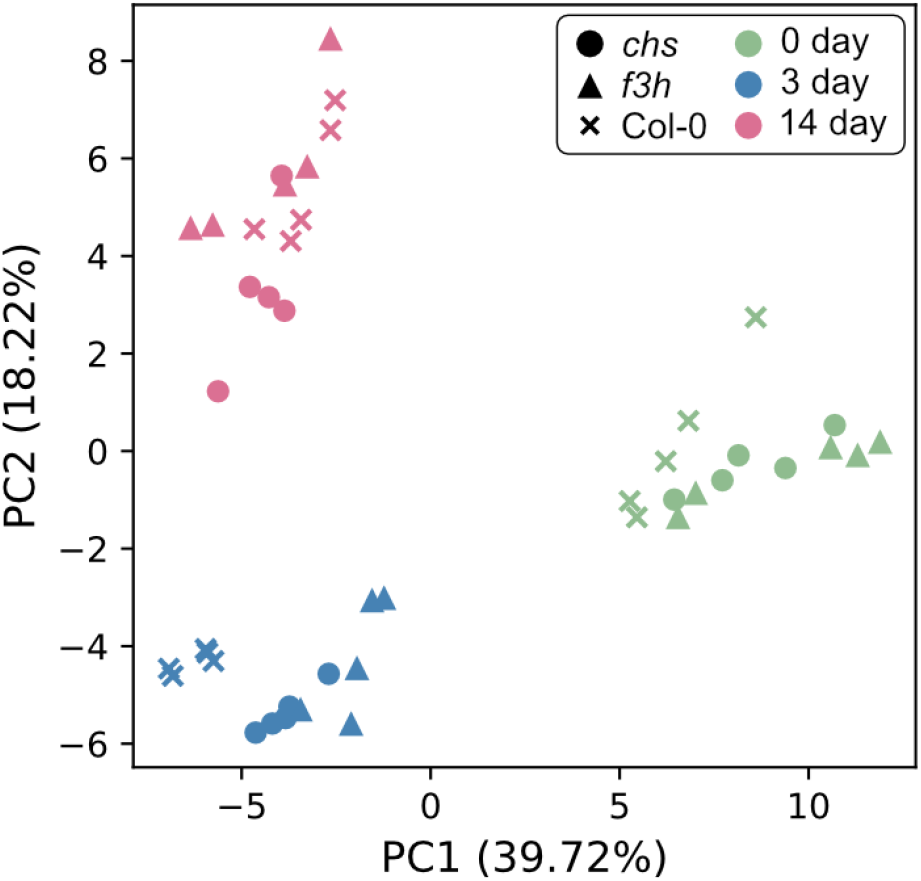
Principal component analysis of the cellular metabolome comprising all considered genotypes and time points of cold acclimation. Circles: *chs*; triangles: *f3h*; crosses: Col-0. Green colour represents: 0 day; blue: 3 day; magenta: 14 days of cold acclimation. Detailed information about loadings and components are provided in the supplements (Supplementary Table 3).

As metabolic acclimation has been found to occur compartment-specific (Nägele and Heyer 2013), subcellular metabolic effects of flavonoid deficiency on primary metabolism were analysed combining non-aqueous fractionation (NAF) of leaf tissue with GC-MS analysis. Comparing subcellular amounts of metabolites before and after cold exposure (0➔3 days, 3➔14 days) revealed that the strongest effects of flavonoid deficiency on cold-induced metabolic reprogramming, which were conserved across *chs* and *f3h* mutants, were located to the plastid and occurred during the early acclimation period, i.e., between 0 and 3 days at 4°C (Figure 4A, Supplementary Figure 5A, B). The metabolites that contributed most to the separation of the *chs* and *f3h* cluster from Col-0 at this time point included proline, glycine, serine, fumarate, aspartate, and malate. Due to their central role in photorespiration, the observed accumulation of glycine and decrease of serine amounts suggested an effect in the regulation of photorespiration in both mutants (Figure 4B). Moreover, glutamine and fumaric acid were found to increase in both mutants while amounts decreased in Col-0 (Figure 4A). Additionally, plants of *chs* were found to accumulate plastidial carbohydrates more than Col-0. These were glucose, fructose, raffinose, glucose 6-phosphate, trehalose, and sucrose after 3 days of cold exposure (Figure 4A).

**Figure 4.**
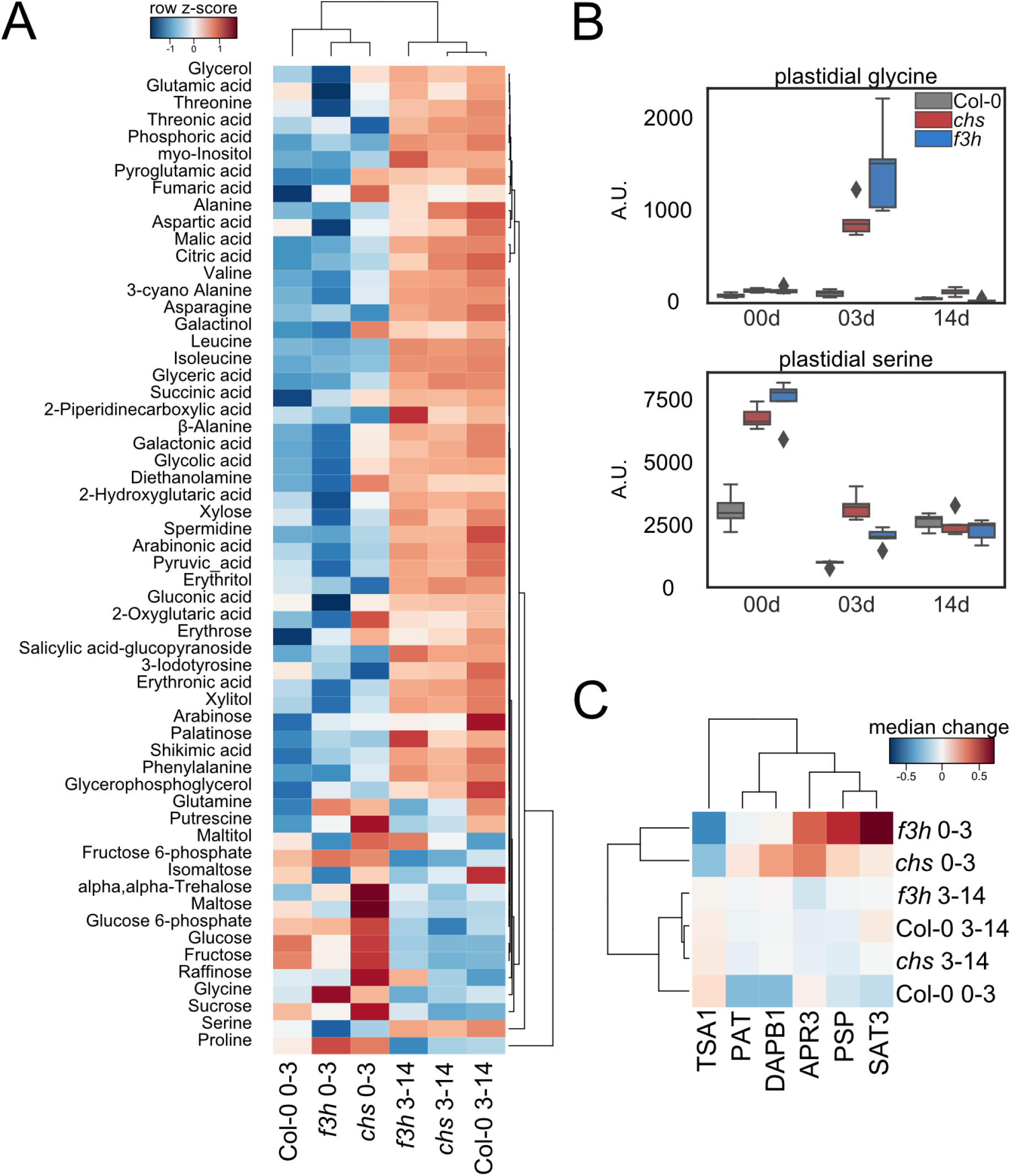
Metabolic cold acclimation in chloroplasts. (**A**) Hierarchical cluster analysis of the subcellular metabolite abundance change rate in plastid between Col-0 and flavonoid mutants. Rates were calculated between 0 and 3 (0-3) and 3 and 14 (3-14) days of cold acclimation (e.g. rate = (median_3 day_ – median_0 day_)/3) in Col-0, *chs*, and *f3h*. Results are displayed in a dendrogram and a heatmap indicating relationships between metabolites and samples based on Euclidean distance and complete-linkage clustering. The values were column-wise standardised using the z-score method. (**B**) Plastidial glycine and serine dynamics in Col-0, *chs*, and *f3h* after 0, 3, and 14 days of cold acclimation. Grey: Col-0; red: *chs*, blue: *f3h*; n=5. The list of significance levels (ANOVA with Tukey HSD) is provided in Supplementary Table 2B. (**C**) Hierarchical cluster analysis of the top 6 amino acid biosynthesis pathway associated protein abundances contributing to the separation of *chs* and *f3h* mutants and Col-0 after 3 days of cold acclimation. Complete information about Euclidean distances in the cluster analysis of in the amino acid biosynthesis related protein dynamics can be found in Supplementary Table 4. Protein abbreviations: APR3 - 5’-adenylylsulfate reductase 3; DAPB1 - 4-hydroxy-tetrahydrodipicolinate reductase 1; PAT - bifunctional aspartate aminotransferase and glutamate/aspartate-prephenate aminotransferase; PSP - phosphoserine phosphatase; SAT3 - serine O-acetyltransferase 3; TSA1 - TSK-associating protein 1.

After 3 days of cold exposure, both mutants showed significantly higher amounts of amino acids than Col-0 in all three analysed compartments (Supplementary Figure 6). In the cytosol and vacuole, there were notable differences between the *chs* and *f3h* mutants. The *f3h* mutant formed a distinct cluster and showed stronger accumulation of amino acids than *chs*. While serine levels did not change significantly in these compartments, glycine peaked after 3 days, especially in the vacuole of both *chs* and *f3h* mutants. Interestingly, both *chs* and *f3h* showed stronger accumulation of cytosolic raffinose (Supplementary Figure 5A).

Given the pronounced effect on amino acids, changes in the proteome were quantified with functional relation to amino acid biosynthesis (Figure 4C, Supplementary Table 4). This showed that, also in the proteome, a similar hierarchical clustering of *chs* and *f3h* mutants became evident after 3 days of cold exposure. The proteins contributing most to this clustering were 5’-adenylylsulfate reductase 3 (APR3), 4-hydroxy-tetrahydrodipicolinate reductase 1 (DAPB1), bifunctional aspartate aminotransferase and glutamate/aspartate-prephenate aminotransferase (PAT), phosphoserine phosphatase (PSP), serine O-acetyltransferase 3 (SAT3), and TSK-associating protein 1 (TSA1; Figure 4C). APR3, PAT, PSP and SAT3 are (in)directly involved in the serine metabolism, which further suggested a differential regulation of the photorespiratory pathway in flavonoid mutants.

Based on the observed differential response of metabolic mutants in amino acids and proteins related to serine metabolism, proteome dynamics in the photorespiratory pathway with its connection to nitrogen, tetrahydrofolate-related C1, sulfur, and methionine metabolism were analysed (Figure 5). The most significant effects in the flavonoid mutants were observed in photorespiration reactions following the serine-hydroxymethyltransferase 1 (SHM1) reaction. In contrast to Col-0 and *f3h*, the *chs* mutant was not significantly affected in SHM1. Simultaneously, the levels of serine:glyoxylate aminotransferase (SGAT), glycerate kinase (GLYK) and plastid glycolate glycerate transporter 1 (PLGG1) decreased in *chs* during the initial 3 days at 4°C. Additionally, ferredoxin-dependent glutamine:oxoglutarate aminotransferase (Fd-GOGAT) and the exchange of glutamic acid and 2-oxoglutaric acid across the plastid membrane were downregulated in *chs* plants. Notably, *chs* mutants showed higher abundances of SHM1 and serine O-acetyltransferase 3 (SAT3), along with slightly elevated abundances of O-acetylserine (thiol) lyase C (OASC). Consequently, dynamics were observed in tetrahydrofolate-related (THF-related) C1 metabolism. 5-formyl-THF acts as a (potential) inhibitor of SHM1, and increased abundance of 5-formyltetrahydrofolate cycloligase (5-FCL), that interconverts 5-formyl-THF into 5,10-methenyl-THF, in *chs* could possibly indicate a feedback regulation of glycine/serine ratio in mitochondria. The plants of *chs* mutants showed an upregulated phosphorylated pathway of serine biosynthesis, a possible response in the plastid to alternations in serine-consuming pathways in the mitochondria. THF-related C1 metabolism connects photorespiration with methionine metabolism in the cytosol, and S-adenosylmethionine synthase (SAM1) abundance showed stronger reduction in Col-0 than in *chs* and *f3h* mutants after 3 days of cold exposure (Figure 5). In summary, cytosolic flavonoid biosynthesis appears crucial for stabilising photorespiration and, consequently, plastidial amino acid metabolism after 3 days of cold exposure.

**Figure 5.**
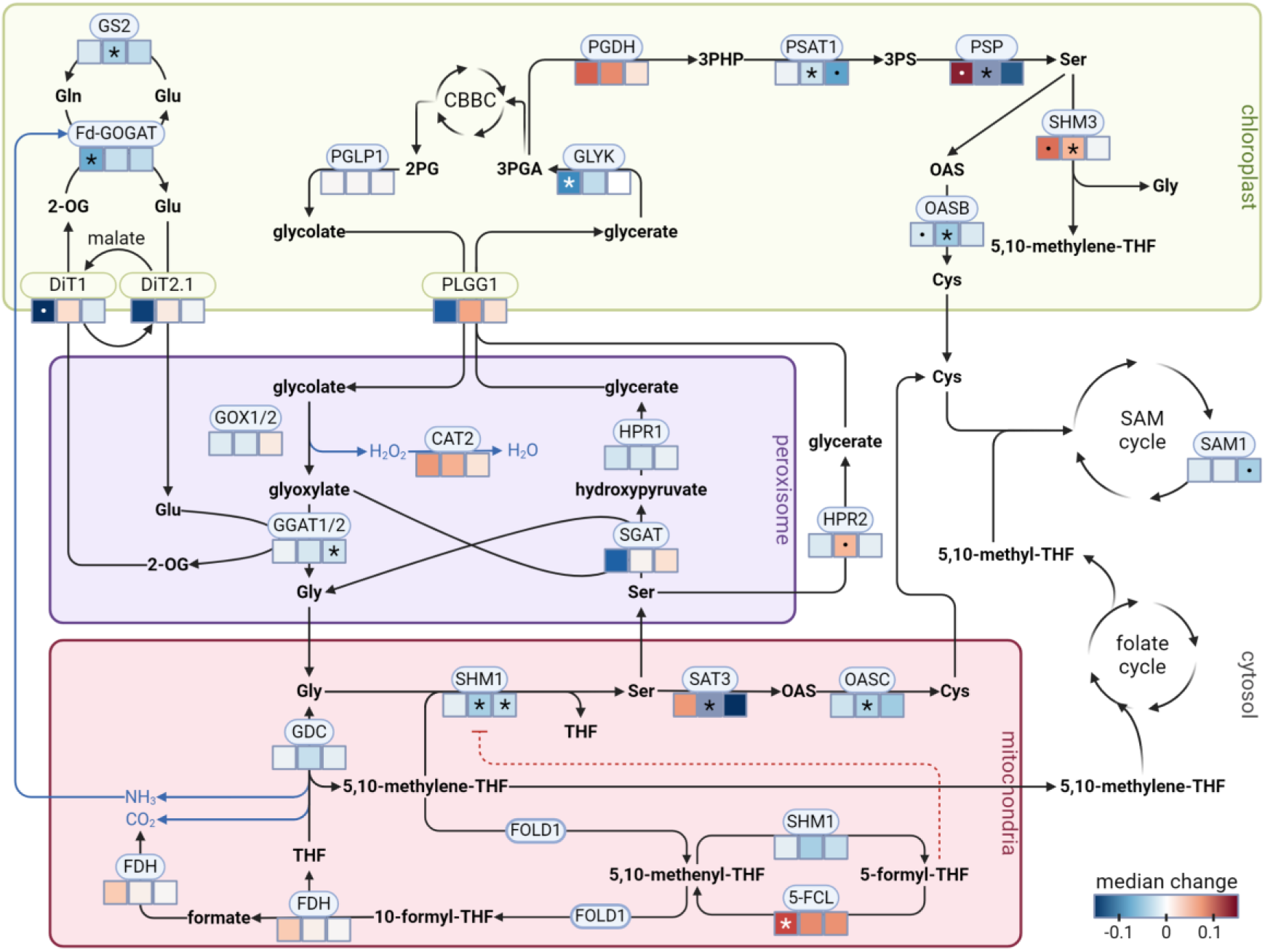
Photorespiration and its interaction with nitrogen, tetrahydrofolate-related C1 and sulfur metabolism. Colour boxes below indicated proteins represent change in average abundance between 0 and 3 days of cold acclimation in *chs* (first box), *f3h* (middle box), and Col-0 (right box). Blue colour represents decrease, red colour represents increase in protein abundance, grey colour for PSP and SAT3 in *f3h* indicates an increase in protein abundance by more than 0.5 A.U. Asterisks and dots indicate significance (t-test; * p < 0.05; **·** p < 0.07. In case of the proteins with several isoforms present, the average median change was calculated. Metabolite abbreviations: 2-OG – 2-oxoglutaric acid; 2PG – 2-phosphoglycolate; 3PGA – 3-phosphoglycerate; 3PHP – 3-phosphohydroxypyruvic acid; 3PS – 3-phosphoserine; Cys – cysteine; Gly – glycine; Gln – glutamine; Glu – glutamic acid; OAS – O-acetylserine; SAM - S-adenosylmethionine; Ser – serine; THF – tetrahydrofolate. Protein abbreviations: 5-FCL - 5-formyltetrahydrofolate cycloligase; CAT2 - catalase 2; DiT1 and DiT2.1 - dicarboxylate transporters; GDC - glycine decarboxylase; GGAT1/2 - glutamate:glyoxylate aminotransferase 1/2; GLYK - glycerate kinase; GOX1/2 - glycolate oxidase 1/2; GS2 - glutamine synthetase 2; HPR1 - hydroxypyruvate reductase 1; HPR2 - hydroxypyruvate reductase 2; Fd-GOGAT - ferredoxin-dependent glutamine:oxoglutarate aminotransferase; FDH - formate dehydrogenase; FOLD1 - bifunctional 5,10-methylene-THF dehydrogenase/5,10-methenyl-THF cyclohydrolase; OASB – O-acetylserine (thiol) lyase B; OASC – O-acetylserine (thiol) lyase C; PGLP1 - phosphoglycolate phosphatase 1; PGDH – 3-phosphoglycerate dehydrogenase; PLGG1 - plastid glycolate glycerate transporter 1; PSAT1 – 3-phosphoserine aminotransferase 1; PSP - phosphoserine phosphatase; SAM1 - S-adenosylmethionine synthase 1; SAT3 - serine O-acetyltransferase 3; SGAT - serine:glyoxylate aminotransferase; SHM1 - serine-hydroxymethyltransferase 1; SHM3 - serine-hydroxymethyltransferase 3.

The absence of flavonoids had impact on both mitochondrial proteins involved in THF-related C1 metabolism, that were detected in the present study, such as 5-formyltetrahydrofolate cycloligase (5-FCL) and formate dehydrogenase (FDH; Figure 5). The THF-related C1 metabolism is connected to the flavonoid biosynthesis pathway through the shikimate pathway, chorismate, and the biosynthesis of para-aminobenzoic acid (Kołton et al. 2022). Although shotgun proteomics applied in the present study could not detect or quantify all proteins of THF metabolism, similar dynamics in aminodeoxychorismate lyase abundance in *chs* and Col-0 suggested that flavonoids might interact with THF-related C1 metabolism not only by influencing precursor availability, but also by direct interactions with central enzymes of this pathway. Evidence from a study on mice suggested a possible direct interaction between isoquercetin and cytosolic C-1-THF synthase (C1TC), a homologue of the bifunctional 5,10-methylene-THF dehydrogenase/5,10-methenyl-THF cyclohydrolase (FOLD1, 2, 3, and 4) proteins in Arabidopsis (Manzoor et al. 2022). The FOLD1 protein and its homologues in plants, yeast, mice, and humans are evolutionarily closely related (Figure 6A). The structural alignment between the FOLD1 protein of *Arabidopsis* and cytosolic C-1-THF synthase of mice showed the root mean square deviation (RMSD) of 1.76 Å, despite only 67% similarity in their amino acid sequences. This indicates that the dehydrogenase/cyclohydrolase region maintains a stable structure while allowing the diverse amino acid combinations (Figure 6B, C). Following this, isoquercetin was identified as a potential ligand for the highly conserved region of FOLD1 protein, forming polar connections with several amino acids (Figure 6D, Supplementary File 1). The negative binding affinity score of -8.79 kcal mol^-1^ indicated an interaction between the protein and the ligand in the identified binding pocket (Guntero et al. 2021; Sefika Feyza et al. 2022). Validation of the molecular docking experiment yielded a binding affinity of -8.94 kcal mol^-1^, with RMSD of 1.49 Å (Supplementary Table 5). In addition to affinity score, negative intermolecular interaction energies, i.e. electrostatic energy and van der Waals energy, of isoquercetin and FOLD1 further supported a possible intermolecular interaction (Supplementary Table 5). These findings suggest a potential role of flavonoids and their precursors in regulating THF-related C1 metabolism, and consequently, photorespiration.

**Figure 6.**
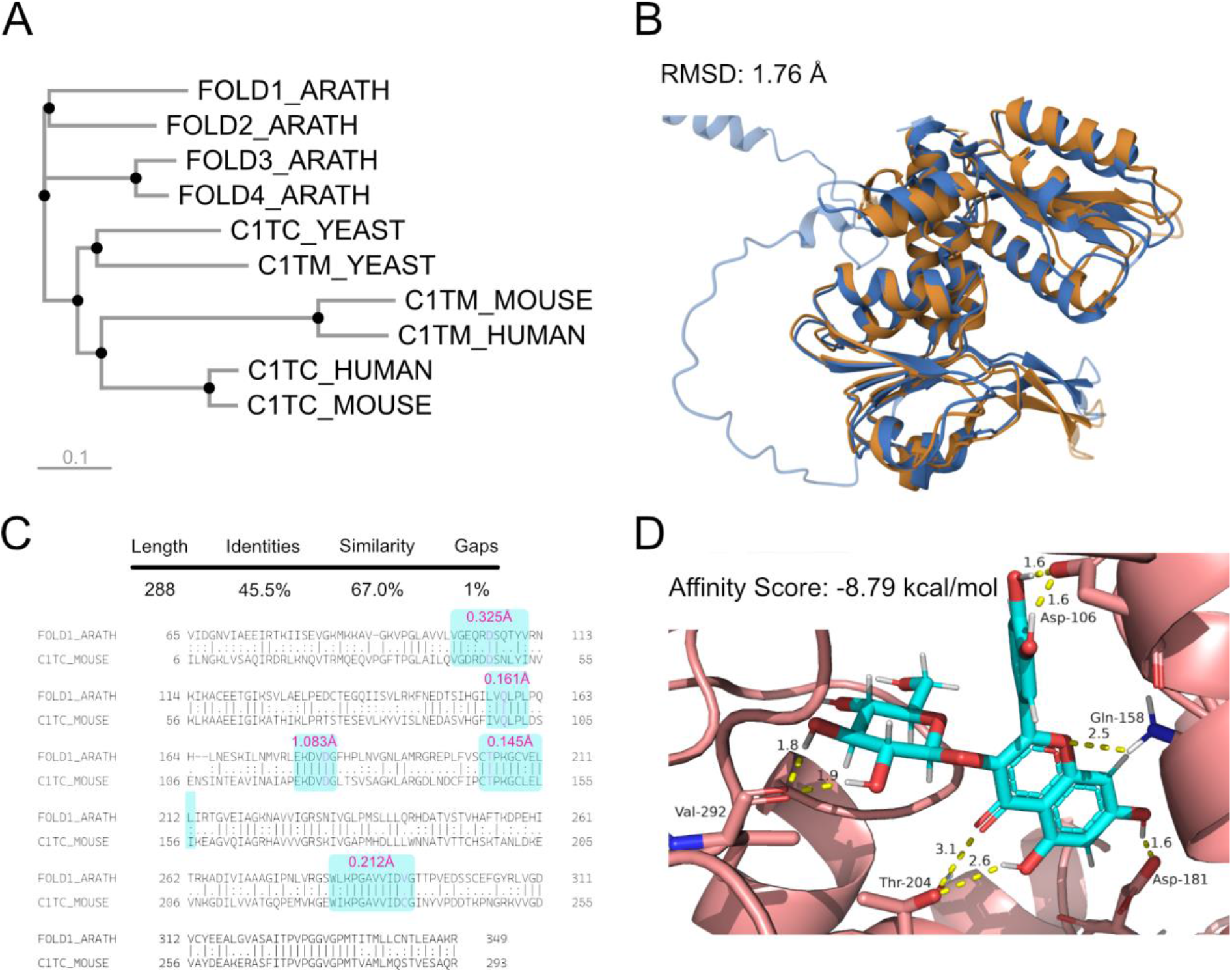
Bifunctional 5,10-methylene-THF dehydrogenase/5,10-methenyl-THF cyclohydrolase (FOLD1) protein analysis. (**A**) Phylogenetic tree of FOLD1 and its homologues: FOLD3/4 – plastidial enzymes of *Arabidopsis*; FOLD2 – cytosolic enzyme of *Arabidopsis*; C1TC – cytosolic enzyme of yeast, mouse and human; C1TM – mitochondrial enzyme of yeast, mouse and human. (**B**) Pairwise protein structure alignment of *Arabidopsis* FOLD1 (blue) and dehydrogenase/cyclohydrolase functional domain of cytosolic mouse C1-THF-synthase protein (sand) with root mean square deviation (RMSD) 1.76 Å. (**C**) Pairwise amino acid sequence alignment of *Arabidopsis* FOLD1 (blue) and dehydrogenase/cyclohydrolase functional domain of cytosolic mouse C1-THF-synthase protein (sand). Turquoise colour indicates highly conservative regions of the protein that potentially interact with isoquercetin molecule. Pink colour indicates interacting with isoquercetin amino acid residues; pink text indicates alignment RMSD of the highlighted regions. (**D**) 3D interaction between the FOLD1 and isoquercetin after molecular docking analysis. Cyan colour: carbon backbone of isoquercetin molecule; salmon: carbon backbone of FOLD1; red: oxygen; grey: hydrogen; blue: nitrogen atoms. Yellow dashed lines represent polar contacts within 5 Å of the isoquercetin molecule, numbers represent distance in Å between the interacting atoms of isoquercetin and amino acid residues. The 3D model is provided in the Supplementary File 1 and the molecular docking energy information is provided in Supplementary Table 5.

Four amino acids involved in the interaction of isoquercetin and FOLD1 (Asp-106, Gln-158, Asp-181 and Thr-204) were identical between *Arabidopsis* and mouse (Figure 6C). A molecular docking experiment on mouse C1TC protein revealed a binding pocket similar to that in *Arabidopsis*, with an affinity score of -8.075 kcal mol^-1^ (Supplementary File 2). This interaction in the C1TC protein involved four amino acids, three of which were identical to Asp-106, Gln-158, and Asp-181 in *Arabidopsis*. Alignment of the individual regions of FOLD1 and C1TC proteins that contain interacting amino acid residues revealed similar 3D structures, with RMSD varying between 0.145 and 1.083 Å. The evolutionary conservation profile of FOLD1 protein revealed these interacting amino acids to be highly conserved and exposed functional residues (Supplementary Figure 7), further supporting the hypothesis of a conserved interaction mechanism between isoquercetin and FOLD1 and its homologues, suggesting a direct regulatory link of flavonoid biosynthesis and THF-related C1 metabolism.

### Protein amounts and regulation of proteasomal compounds are associated with flavonoid biosynthesis

Deficiencies in flavonoid biosynthesis were found to affect amino acid metabolism (Figure 1, Figure 4, Figure 5). To determine whether these alternations in amino acid metabolism might also have an impact on protein homeostasis, the total protein amount was quantified and normalised to dry weight (Figure 7A). During the first 3 days of cold acclimation, the total protein amount decreased across all genotypes, with the *chs* and *f3h* mutants showing a more pronounced decrease than Col-0. Additionally, between 3 and 14 days, protein amounts in both mutants were found to further decline, while Col-0 stabilised its protein homeostasis. Overall, between 0 and 14 days, the *chs* mutant showed the most significant reduction in total protein level, with a median decrease of more than 2-fold. In contrast, Col-0 showed a stabilised protein content after 14 days of cold acclimation, with only 1.2-fold decrease (Figure 7A). After 3 days of cold, both *chs* and *f3h* mutants exhibited similar or higher levels of amino acids in plastid, cytosol and vacuole compared to Col-0 (Supplementary Figure 6). After 14 days, both mutants showed reduced amino acid levels in the cytosol, although *chs* still had more than after 3 days. Only in *f3h*, amino acid levels in the cytosol significantly decreased after 14 days, indicating that the protein decrease in *chs* and *f3h* was not due to a general lack of substrates for protein biosynthesis. The minor decline in amino acids was observed only in *f3h* in cytosol (Supplementary Figure 6), suggesting alterations in the regulation of the protein biosynthesis and/or degradation machinery. Quantification of cytosolic ribosomal proteins, which mediate the biosynthesis of most proteins in plants cell and are often associated to the rough endoplasmic reticulum (ER), showed an insignificant differential trend in *chs* plants, with ribosome abundance slightly decreasing during the first 3 days at low temperature (Figure 7B). In contrast, Col-0 and *f3h* medians increased in this period. On the other hand, during the same period, proteasomal protein abundances, responsible for protein degradation and also associated to the ER, decreased in Col-0 but increased in both flavonoid mutants, potentially contributing to the initial decline in protein content (Figure 7C).

**Figure 7.**
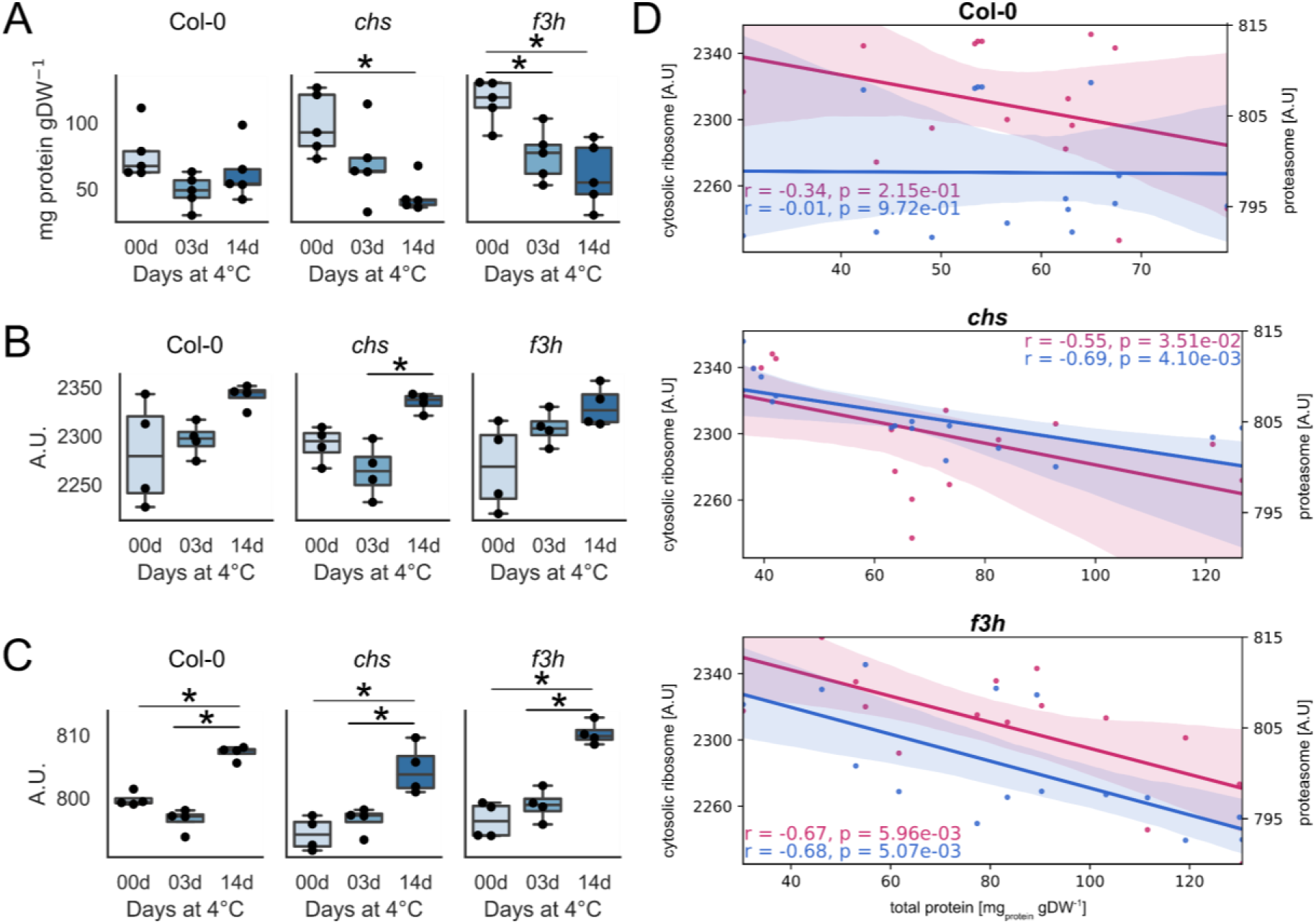
Regulation of total protein amount during cold acclimation. (**A**) Total protein amount; (**B**) cytosolic ribosome, and (**C**) proteasome associated protein abundance in Col-0, *chs*, and *f3h* before (0 day) and after (3 or 14 days) of cold acclimation. Points represent biological replicates (n≥4), asterisks represent significant differences (ANOVA with Tukey HSD, p-value<0.05), the full list of significance levels is provided in Supplementary Table 2A. (**D**) Pearson correlation analysis between total protein amount (x-axis) and cytosolic ribosome associated protein abundance (primary y-axis, pink), and between total protein amount (x-axis) and proteasome associated protein abundance (secondary y-axis, blue) in Col-0, *chs* and *f3h* across 0, 3 and 14 days of cold acclimation. Points represent independent replicates, n=5; solid line represents linear regression and shaded area represents confidence interval of the linear regression.

After 14 days, despite the opposite trends in total protein levels between Col-0 and the flavonoid mutants, both *chs* and *f3h* demonstrated a wild type-like increase in cytosolic ribosome and proteasome abundance (Figure 7B, C). Pearson correlation analysis between total protein amount and either cytosolic ribosome abundance or proteasome abundance revealed a strong significant negative correlation throughout the whole period of cold acclimation in *chs* and *f3h*, but not in Col-0 (Figure 7D). This finding suggests a complex interaction between ribosome and proteasome machineries and flavonoid accumulation during the later stages of cold acclimation.

Differential protein abundance GO enrichment analysis on the whole proteome dynamics in the *chs* mutant after 14 days of cold revealed upregulation of proteins related to cytoplasmic translation, proteasomal protein catabolic process, and ribosome assembly (Supplementary Table 6). Moreover, unlike Col-0, both *chs* and *f3h* did not show an upregulation of ‘protein folding’. In summary, these results suggest an effect (direct or indirect) of flavonoids on protein biosynthesis, degradation and folding.

## Discussion

Metabolic acclimation of plants during cold exposure has been described in many studies to comprise and affect a large array of enzymatic reactions and transport processes across intracellular membrane systems which ultimately results in adjustment of metabolite concentrations, see e.g. (Hannah et al. 2006; Korn et al. 2008; Koster and Lynch 1992; Hoermiller et al. 2017; Garcia-Molina et al. 2020). Although many of these metabolic adjustments are strong and significant effects, frequently, their physiological function remains elusive. Particularly, this might be due to the high number of variables which contribute to such metabolite dynamics: enzymes involved in biosynthesis, degradation and compartmental sequestration. For example, accumulation of soluble carbohydrates, e.g., sucrose, hexoses and raffinose oligosaccharides (RFOs) is a well-known cold response of diverse plant species (Koster and Lynch 1992; Alberdi et al. 1993; Klotke et al. 2004; Kaplan et al. 2004). Although biophysical studies provided evidence for protective functions of carbohydrates during cold and freezing *in vitro* (Hincha et al. 2003), *in vivo* function can hardly be predicted without subcellular localization (Zuther et al. 2004; Knaupp et al. 2011). Based on such observations, in the present study, plant metabolism was resolved to a subcellular and compartment-specific level to reduce ambiguity of functional interpretation of metabolite dynamics. Time series analysis of GC-MS derived metabolomics data unravelled a cold-induced perturbation of citrate and glutamate metabolism in *chs* and *f3h* mutants (Figure 1, Supplementary Figure 2). Subcellular proteomics suggested a shift of ICDH activity and/or flux from mitochondria to the cytosol due to flavonoid deficiency (Figure 2, Supplementary Figure 3). Such a cytosolic bypass was discussed earlier in context of C/N balance and interconversion of a citrate storage outside the mitochondria (Sweetlove et al. 2010). Even more, for illuminated leaves, such non-cyclic TCA cycle flux modes were shown to reduce PDH-catalysed decarboxylation rates by up to 75% (Tcherkez et al. 2009). Carbon skeletons of such fluxes are then substrate for glutamate and glutamine biosynthesis. Due to the finding that both metabolites over-accumulated in cytosol and plastids of *chs* and *f3h* until 3 days of cold acclimation, this suggests that flavonoid metabolism affects, or even co-regulates, the C/N balance during cold exposure. Previous work suggested that a high C/N ratio under low nitrogen availability increases flavonoid content (Li et al. 2023). Further, flavonoid pathway activators *PAP1* and *PAP2* were found to strongly respond to low temperature and low nitrogen (Olsen et al. 2009). Together with the data of the present study this suggests that C/N balance and flavonoid metabolism are mutually regulated. This is also supported by the finding that in *chs*, both GOGAT enzymes and plastidial dicarboxylate transporters were significantly downregulated during the initial cold acclimation period. Hence, during the early phase of cold acclimation, flavonoid biosynthesis may stabilise plastidial C/N metabolism and, indirectly, photosynthesis by consuming carbon skeletons in the cytosol which originate from citrate metabolism. Finally, this might connect mitochondrial, cytosolic and plastidial redox and energy balance under environmental fluctuation.

Flavonoid biosynthesis results from a combination of phenylpropanoid pathway and the acetate pathway (Perez de Souza et al. 2020). Based on the observation that cICDH positively correlated with pPDH (see Figure 2), this suggests that flavonoid deficiency resulted in an increased flux through the acetate pathway which supplied malonyl-CoA moieties for fatty acid and lipid biosynthesis. This is supported by the finding that triacylglycerols were significantly elevated in both mutants under cold (Supplementary Tables 3 and 7).

The analysis of the primary metabolomes revealed a strong and significant effect of flavonoid deficiency on amino acid metabolism, and this effect was most pronounced in plastids (see Figure 4). While it may not be surprising that metabolism of aromatic amino acids, which are direct substrates for flavonoid biosynthesis, are affected in *chs* and *f3h* mutants, it is even more interesting to see emphasized effects in the photorespiratory intermediates glycine and serine. Due to experimental limitations of the NAF protocol, in which separation of plastidial and mitochondrial metabolomes remains difficult (Fürtauer et al. 2019), the presented data cannot be exclusively interpreted as plastidial or mitochondrial metabolite concentrations. However, due to the accumulation of the substrate glycine and the depletion of the product serine of the mitochondrial SHM1-catalyzed reaction, it appears plausible that the observed metabolic phenotype is due to affected photorespiratory regulation. Further, quantified protein dynamics suggested that deficiencies in the flavonoid biosynthesis pathway resulted in upregulated plastidial serine biosynthesis (see Figure 5). This might be a regulatory consequence of a depleted mitochondrial serine pool which limits its availability for cellular functions and might be compensated for by this plastidial bypass. Plastidial proteins PGDH, PSP and SHM3 were found to be upregulated in both *chs* and *f3h* plants. Simultaneously, cysteine biosynthesis via O-acetylserine was upregulated during the first 3 days of cold acclimation in these mutants which further indicates that mitochondrial deficiencies in serine biosynthesis may have been compensated for in the plastids.

In mitochondria, SHM1 catalyzes the interconversion of glycine to serine which comprises a methylene transfer from 5,10-methylene-THF (Shi and Bloom 2021). Alternatively, 5,10-methylene-THF might be interconverted into 5,10-methenly-THF, catalyzed by FOLD1 (AT2G38660), which represents a bifunctional 5,10-methylene-THF dehydrogenase/5,10-methenyl-THF cyclohydrolase being localized to mitochondria (Collakova et al. 2008; Groth et al. 2016). The competitive inhibitor of SHM1 and other folate-dependent enzymes, 5-formyl-THF is biosynthesized by SHM1 in the presence of glycine (Stover and Schirch 1990). In plants of *chs*, a significant and strong upregulation of 5-FCL during the first 3 days at 4°C was observed which metabolizes 5-formyl-THF (Roje et al. 2002). Although 5-FCL was also upregulated in Col-0, the upregulation was stronger in both flavonoid mutants and only significant in *chs* (Figure 5). Thus, a potential explanation for this observation might be that metabolic compounds or pathways which are affected by the *chs* mutation might represent potential inhibitors of 5-formyl-THF. Indeed, for the FOLD1-homolgue in mice, C1TC, was observed to be inhibited by isoquercetin, which is produced downstream of CHS (Manzoor et al. 2022). Surprisingly, both metabolomic and proteomic dynamics in these pathways were most significantly affected in mutants during the early phase of cold acclimation in which accumulation of anthocyanins, i.e., late products of the flavonoid pathway, in Col-0 was less pronounced than after 14 days (for anthocyanin data, please refer to (Kitashova et al. 2023)). This observation suggests that early intermediates of flavonoid biosynthesis play a crucial role for metabolic cold acclimation but clearly differ in their function from end products of the same pathway which might rather have a function as light or UV protectants (Zirngibl et al. 2023; Zhang et al. 2017; Li et al. 1993).

### Flavonoid metabolism stabilizes protein homeostasis during cold acclimation

Cold acclimation resulted in a decrease of protein amounts normalized to sample dry mass (see Figure 7). While Col-0 was found to successfully stabilize its protein amount on a slightly lower level than at 22°C, both *f3h* and *chs* mutants showed a significant decrease over the full cold acclimation period. Proteomics data revealed that in both flavonoid mutants proteasomal complexes were upregulated during the initial phase of cold acclimation, i.e., until 3 days at 4°C, while a negative trend was observed for Col-0. This resulted in significantly negative correlations of proteasome and protein amounts in *f3h* and *chs* while no significant correlation was observed in Col-0. This points towards a function of flavonoids and/or intermediates of flavonoid biosynthesis in stabilization of the protein homeostasis at low temperature.

Interestingly, in a GO term enrichment analysis, ‘protein folding’ was significantly over-represented in *chs* between 0 and 3 days at 4°C while no significance was detected in Col-0 (Supplementary Table 6). The functional category ‘protein refolding’ was decreased in this period in Col-0 while it remained constant in both mutants. While, to the best of our knowledge, up to today there is no direct experimental proof for flavonoid-based stabilization of proteins during plant cold acclimation, such interaction has been shown in other systems. For example, stability and folding of the visual G protein-coupled receptor rhodopsin, which is expressed in the rod photoreceptors of the eye, was shown to be enhanced by flavonoids, e.g., quercetin or myricetin (Ortega et al. 2019). It was observed that flavonoids modulate the protein’s conformation by direct interaction and stabilise it, probably by introducing structural rigidity. Hence, deficiencies of flavonoids might lead in misfolded proteins which subsequently undergo refolding or proteasomal degradation. Another study showed that flavonoids inhibited proteasome 26S activity in pig red blood cells (Chang 2009).

Although the effect on ribosomes in *chs* and *f3h* was less pronounced than on the proteasome, it still became significant in the correlation with protein amounts (see Figure 7D). Previous work has highlighted the important role of ribosomes and translational regulation during cold acclimation (Garcia-Molina et al. 2020). In line with this, data of the present study suggests that flavonoids influence protein homeostasis during cold exposure, maybe through their direct interactions with ribosomes or tRNA (Kanakis et al. 2006). In summary, although it remains speculation from experimental data of the present study, it seems plausible that also in plants, such interactions between flavonoids and proteins as well as the proteasome, ribosomes and/or tRNAs might occur and contribute to a stable protein homeostasis at low temperature.

## Conclusion

Findings of the present study indicate that flavonoids play different physiological roles during a two-week cold acclimation period. During the initial cold response, flavonoid biosynthesis was found to affect, and maybe regulate, stabilisation of the C/N balance between plastids, cytosol, and mitochondria. During the late cold acclimation period, flavonoids affect protein stability and folding which has dramatic effects on a plant’s energy homeostasis. It remains to be elucidated whether such functional diversity can also be observed under different flavonoid-stimulating conditions, such as high light or excess UV. Finally, however, the physiological function of flavonoids and involved pathways clearly goes beyond the accumulation of pigments for dissipation of electromagnetic radiation energy.

## Acknowledgements

We thank the members of Plant Evolutionary Cell Biology at LMU München and the members of TRR175 for constructive discussions and advice. Further, we thank the team of MSBioLMU as well as the Graduate School Life Science Munich (LSM) for support. This work was funded by Deutsche Forschungsgemeinschaft (DFG), TRR175/D03 and TRR175/Z1.

## Author Contributions

A.K. performed fractionation experiments, statistical analysis and data evaluation, and wrote the paper. M.L. performed metabolomics analyses, S.S. and D.L. performed, supervised and conceived proteomics analyses; T.N. conceived the study, performed statistics and data evaluation and wrote the paper.

## Supplementary Figures

**Supplementary Figure 1.**
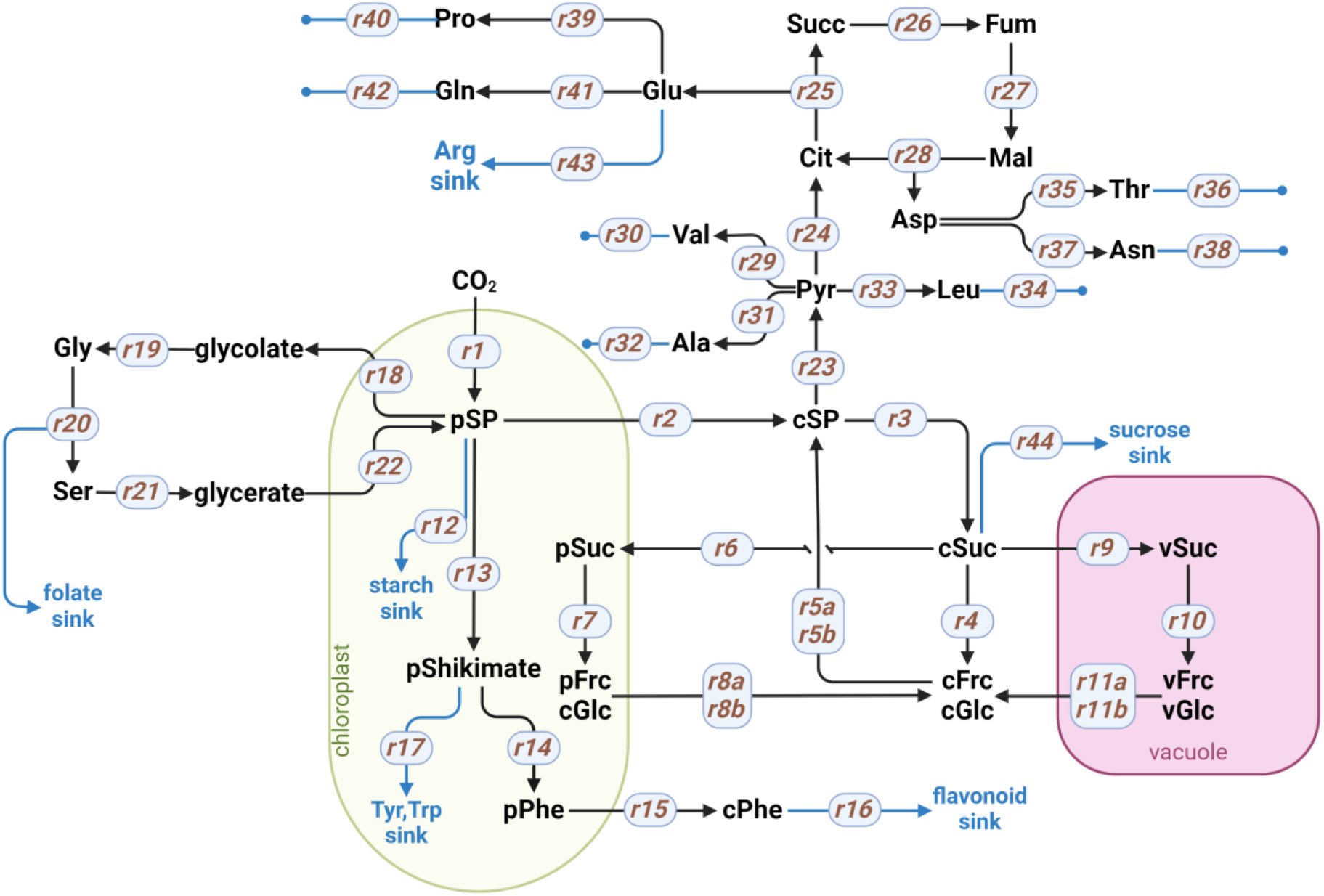
Graphical representation of the metabolic network of subcellular carbohydrate metabolism, carboxylic acids and amino acids used for the analysis of the dynamics of subcellular metabolomes. Ala: alanine; Arg: arginine; Asn: asparagine; Asp: aspartic acid; Cit: citric acid; cFrc: cytosolic fructose; pFrc: plastidial fructose; vFrc: vacuolar fructose; Fum: fumaric acid; cGlc: cytosolic glucose; pGlc: plastidial glucose; vGlc: vacuolar glucose; Gln: glutamine; Glu: glutamic acid; Gly: glycine; Leu: leucine; Mal: malic acid; cPhe: cytosolic phenylalanine; pPhe: plastidial phenylalanine; Pro: proline; Pyr: pyruvic acid; Thr: threonine; Ser: serine; cSP: cytosolic sugar phosphates; pSP: plastidial sugar phosphates; cSuc: cytosolic sucrose; pSuc: plastidial sucrose; vSuc: vacuolar sucrose; Succ: succinic acid; Trp: tryptophan; Tyr: tyrosine; Val: valine.

**Supplementary Figure 2.**
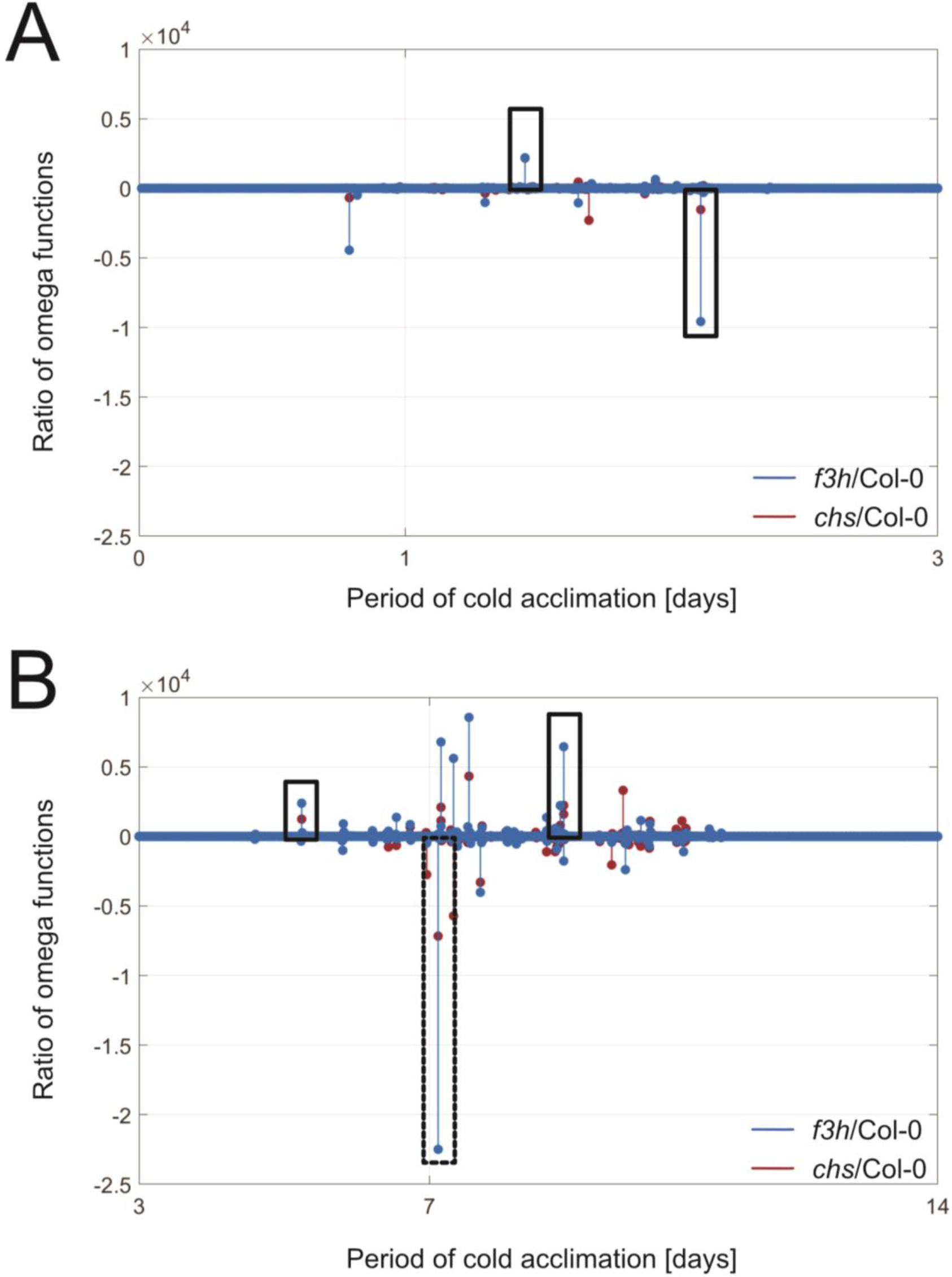
Ratios of omega functions revealed by time series analysis. Omega functions reflected changes in dynamics between substrate and product amounts of enzyme reactions. **(A)** Scaled ratios of omega functions during the cold acclimation period 0➔3 days. Red: ratio *f3h*/Col-0; blue: ratio *chs*/Col-0. **(B)** Scaled ratios of omega functions during the cold acclimation period 3➔14 days. Red: ratio *f3h*/Col-0; blue: ratio *chs*/Col-0. Detailed information about ratios is provided in Supplementary Table 1. Black rectangles indicate conserved metabolic interactions of citrate and glutamate metabolism which were found to be perturbed consistently in both mutants and across the full acclimation period. Black rectangle with dashed lines indicates the strongest deflection in both mutants which was a deregulated citrate interconversion.

**Supplementary Figure 3.**
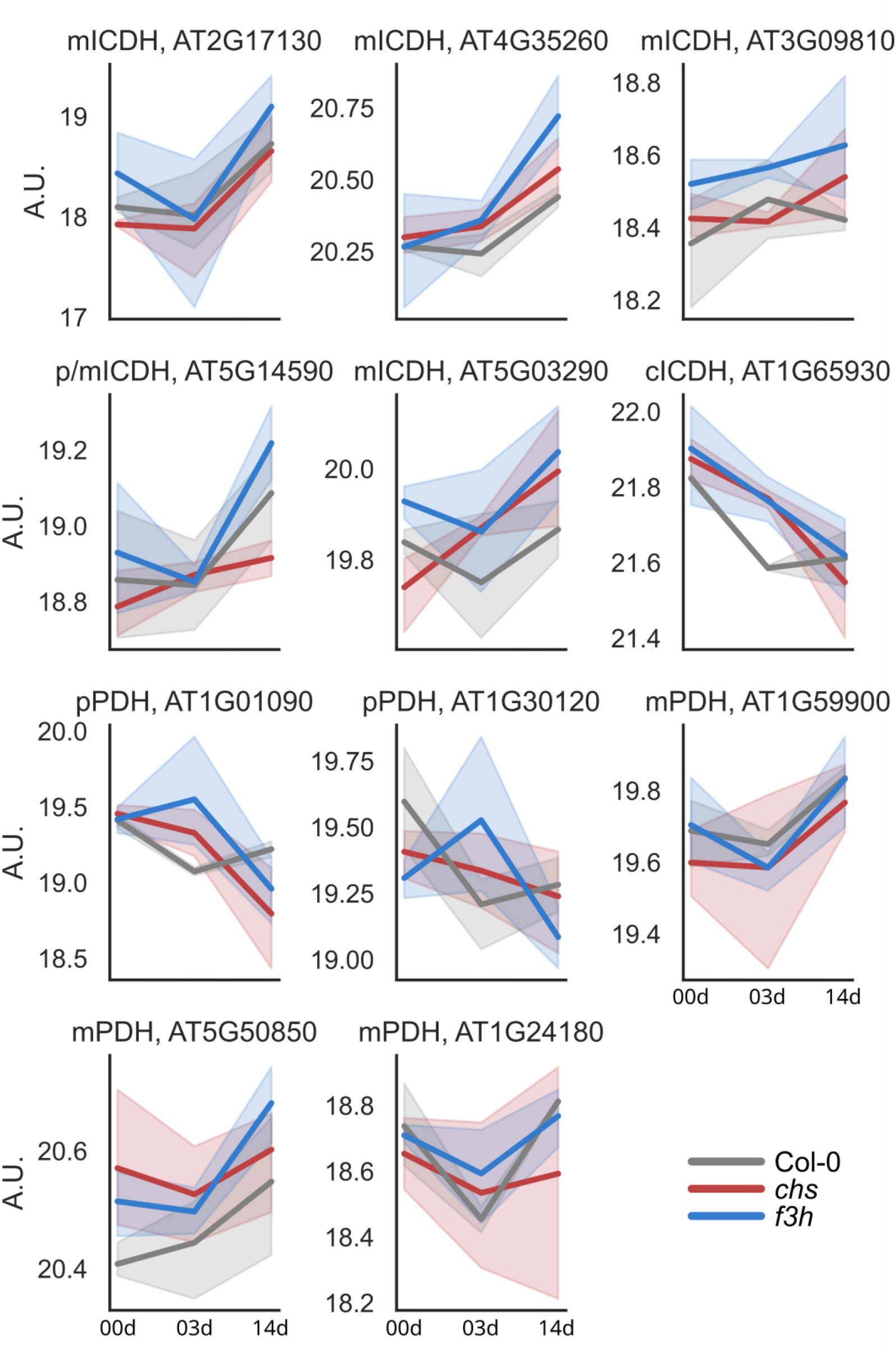
Protein abundance dynamics in Col-0, *chs* and *f3h* after 0, 3 and 14 days of cold acclimation. mICDH: mitochondrial isocitrate dehydrogenase; p/mICDH: plastidial/mitochondrial isocitrate dehydrogenase; cICDH: cytosolic isocitrate dehydrogenase; pPDH: plastidial pyruvate dehydrogenase; mPDH: mitochondrial pyruvate dehydrogenase. Solid line represents change of mean abundance; and shaded area represents standard deviation; n=4. Grey: Col-0; red: *chs*; blue: *f3h*.

**Supplementary Figure 4.**
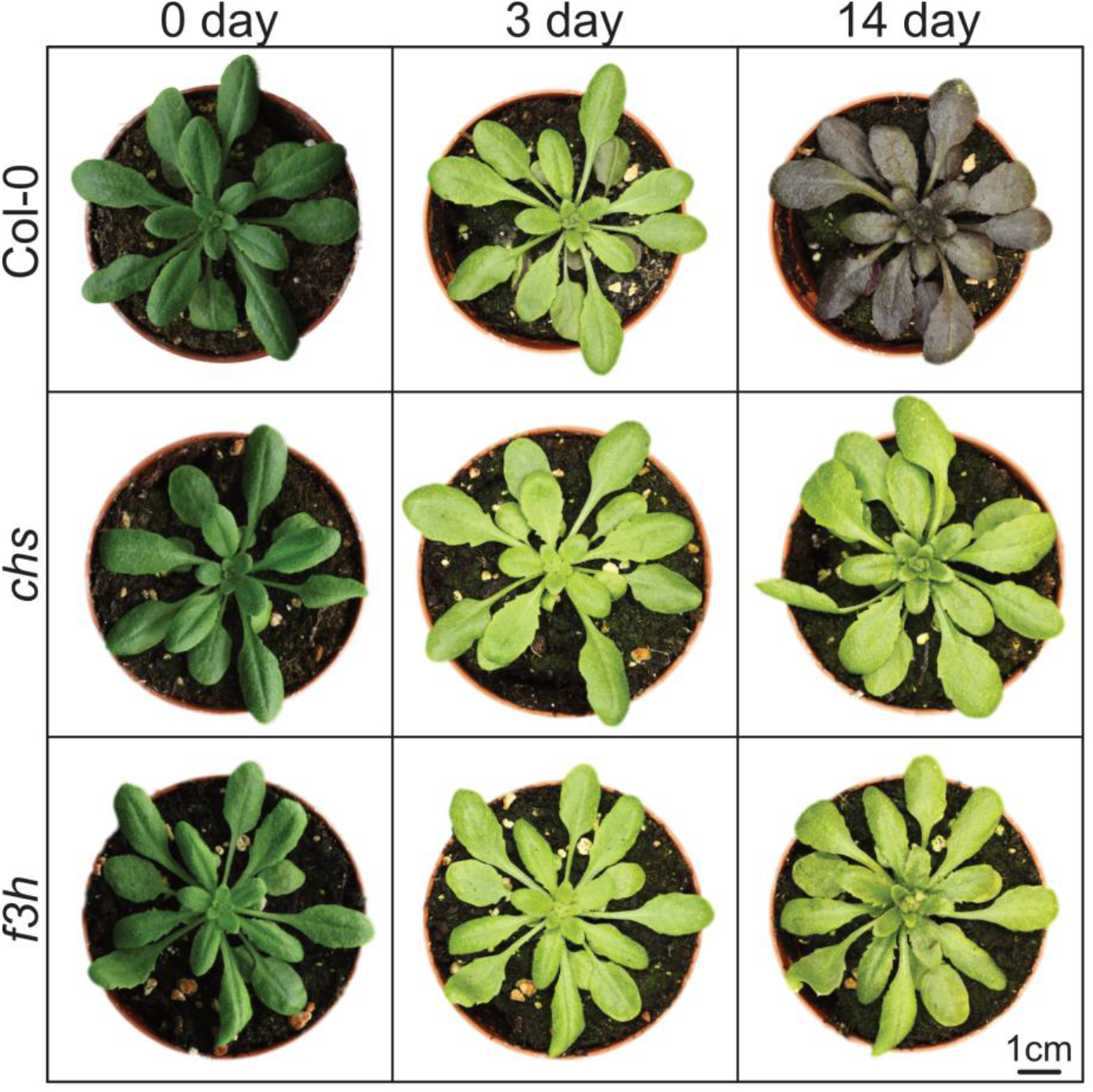
Growth phenotypes of Col-0, *chs* and *f3h* after 0, 3 and 14 days of cold acclimation. Quantified amounts of anthocyanins were provided earlier (Kitashova et al. 2023). Variation in the shades of green between non-acclimated and acclimated plants were due to different illumination levels during imaging.

**Supplementary Figure 5.**
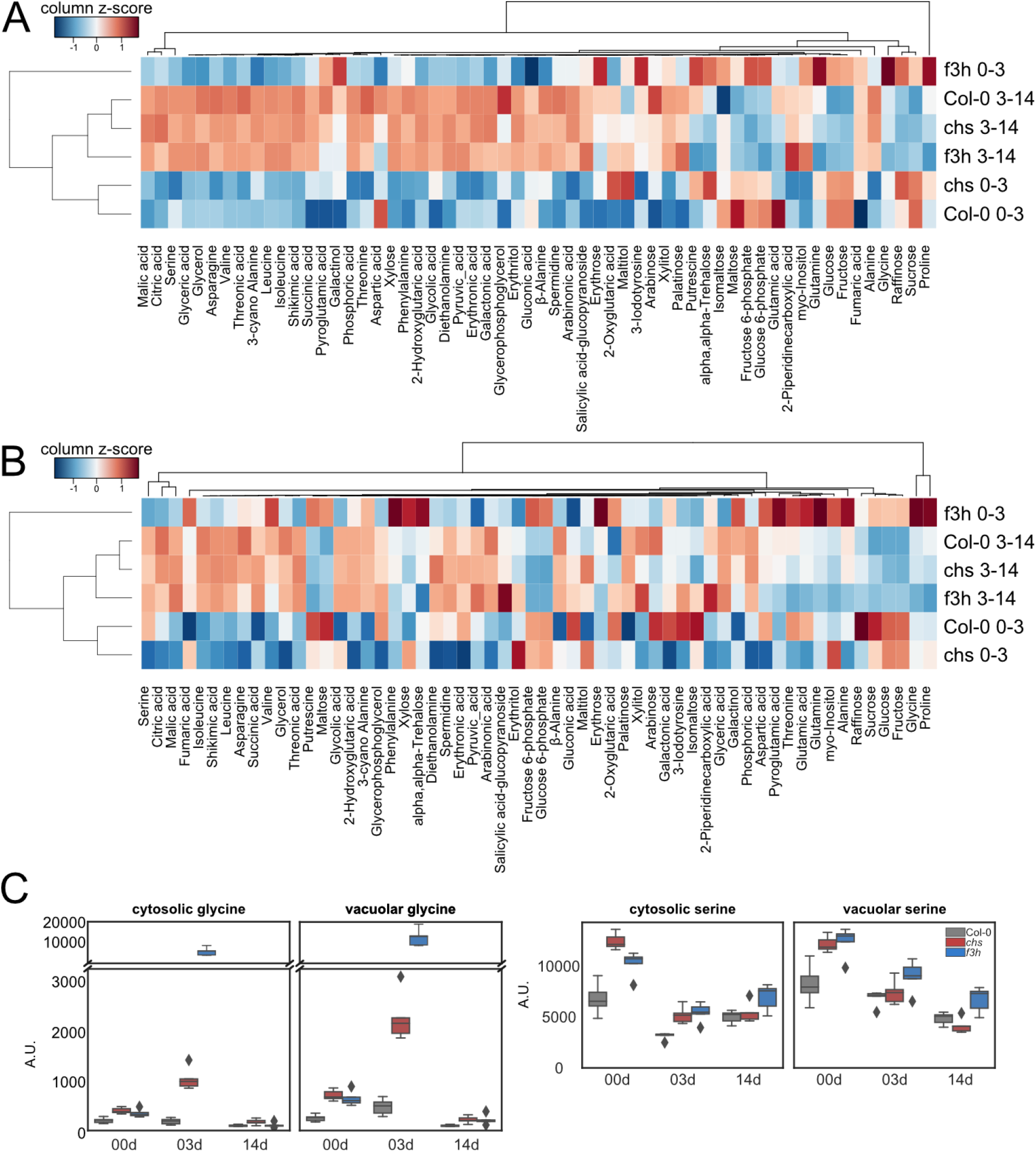
Hierarchical cluster analysis of the subcellular metabolite abundance change rate. **(A)** cytosol; and **(B)** vacuole between 0 and 3 (0-3) and 3 and 14 (3-14) days of cold acclimation (e.g. rate = (median_3 day_ – median_0 day_)/3) in Col-0, *chs*, and *f3h*. Results are displayed in a dendrogram and a heatmap indicating relationships between metabolites and samples based on Euclidean distance and complete-linkage clustering. The values were column-wise standardised using the z-score method. **(C)** Glycine and serine dynamics in cytosol and vacuole in Col-0, *chs* and *f3h* after 0, 3, and 14 days of cold acclimation. Grey: Col-0; red: *chs*; blue: *f3h*; n=5. The list of significance levels (ANOVA with Tukey HSD) is provided in Supplementary Table 2B.

**Supplementary Figure 6.**
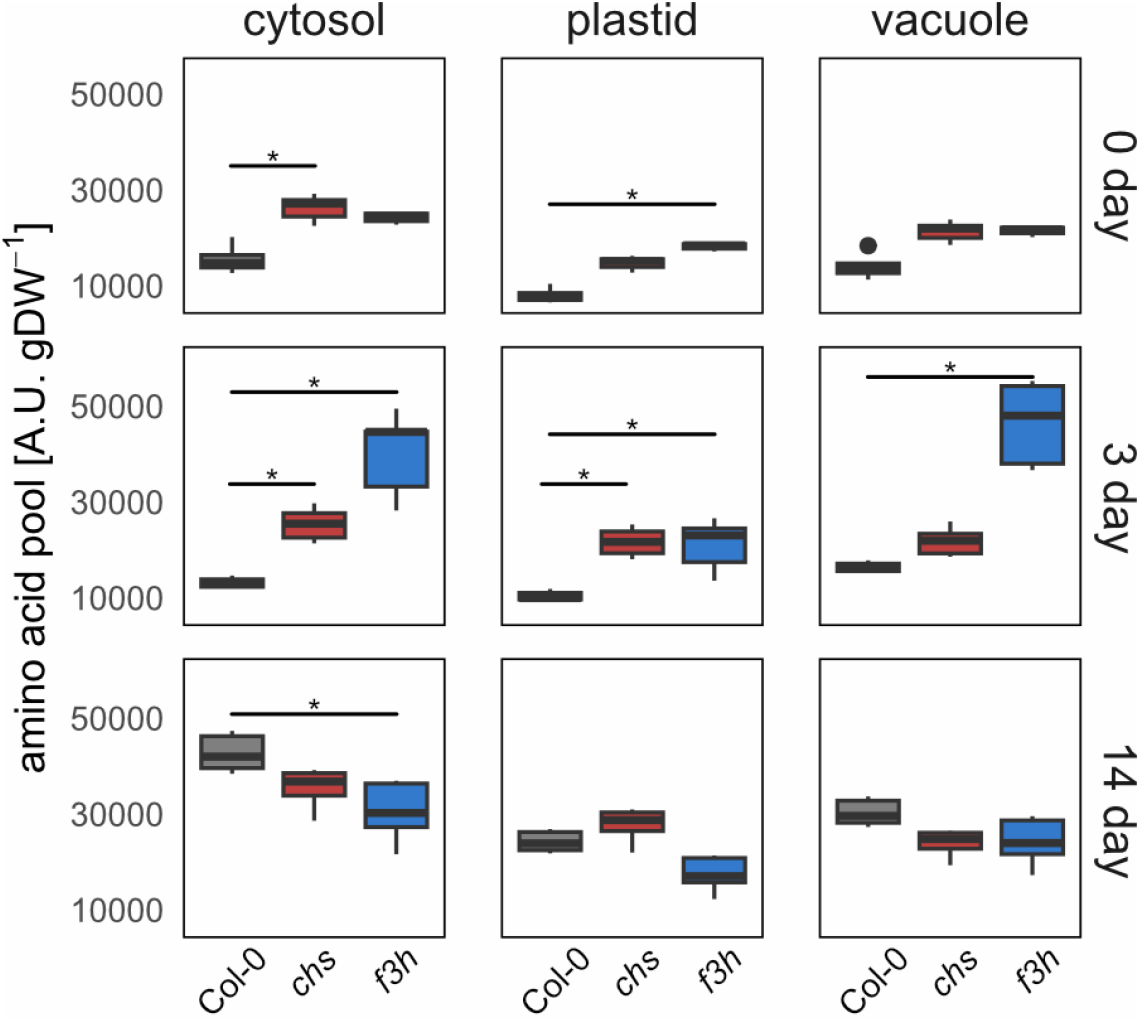
Dynamics of main amino acid pool in Col-0, *chs* and *f3h* in cytosol, plastid and vacuole after 0, 3 and 14 days of cold acclimation. Asterisks indicate significant difference between mutants and wild type (ANOVA with Tukey HSD, p-value < 0.05, n=5). The full list of significance levels is provided in Supplementary Table 2B.

**Supplementary Figure 7.**
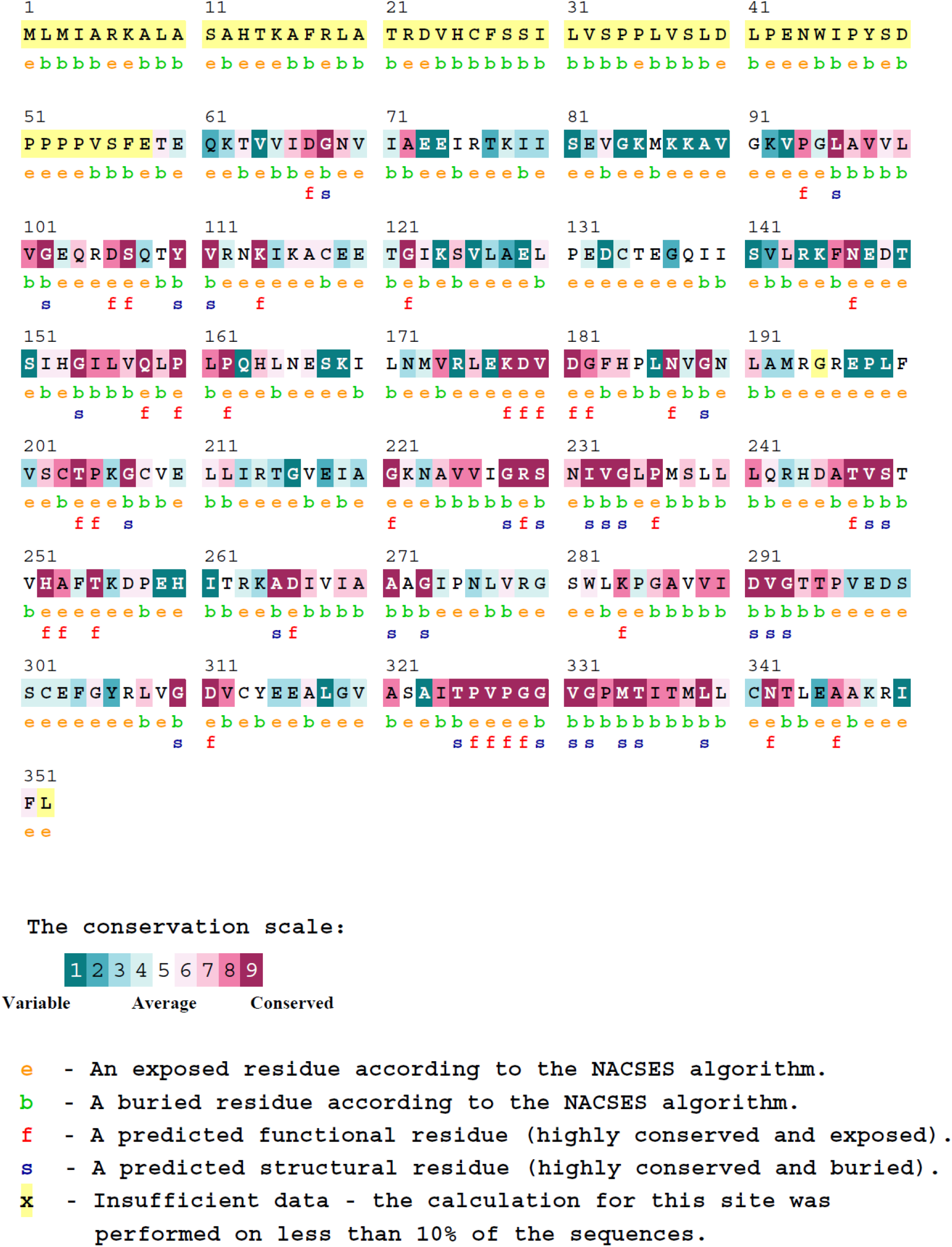
Evolutionary conservation profile of FOLD1 protein. The calculation was performed on 150 sequences that are homologous to the FOLD1 protein with ConSurf software.

## Supplementary legends

**Supplementary Table 1. Ratios of scaled time series omega functions.** Interpolated Gompertz functions were used to calculated omega functions (see Methods section). Functions were scaled (z-scale, i.e., zero mean – unit variance) before ratios of function values were built for (**A**) chs/Col-0 (0➔3d), (**B**) f3h/Col-0 (0➔3d), (**C**) chs/Col-0 (3➔14d), (**D**) f3h/Col-0 (3➔14d).

**Supplementary Table 2. ANOVA with Tukey HSD results**. Significance levels for **(A)** proteome level; **(B)** subcellular metabolomics; and **(C)** cellular metabolomics comparisons, p-values below 0.05 highlighted in green.

**Supplementary Table 3. Principal Component Analysis (PCA) loadings and component scores for metabolites quantified.**

**Supplementary Table 4. Cluster analysis result for proteins involved in amino acid biosynthesis. (A)** List of proteins involved in amino acid biosynthesis; **(B)** row-wise and **(C)** column-wise calculated Euclidean distances.

**Supplementary Table 5. Summary of molecular docking results for isoquercetin with bifunctional 5,10-methylene-THF dehydrogenase/5,10-methenyl-THF cyclohydrolase (FOLD1) protein.**

**Supplementary Table 6. GO term enrichment analysis.** Differentially abundant proteins in chs and f3h were identified based on ANOVA with Tukey HSD results between 0 and 3 days, and 3 and 14 days of cold acclimation (Supplementary Table 2A). Only GO terms related to biological processes are included in the analysis.

**Supplementary Table 7. Cold-induced dynamics of metabolome in Col-0, *chs* and *f3h* after 0, 3 and 14 days of cold acclimation.** Mean values and standard deviations are provided for (A) subcellular) and (B) cellular metabolite amounts.

**Supplementary File 1. 3D structure of isoquercetin and bifunctional 5,10-methylene-THF dehydrogenase/5,10-methenyl-THF cyclohydrolase (FOLD1) protein of *Arabidopsis* interaction.**

**Supplementary File 2. 3D structure of isoquercetin and C-1-THF synthase protein of mouse interaction.**

